# Distinct phases of cellular signaling revealed by time-resolved protein synthesis

**DOI:** 10.1101/2023.07.10.548208

**Authors:** Gihoon Lee, Tom W. Muir

## Abstract

The post-translational regulation of protein function is involved in most cellular processes. As such, synthetic biology tools that operate at this level provide opportunities for manipulating cellular states. Here, we deploy a proximity-triggered protein trans-splicing technology to enable the time-resolved synthesis of target proteins from pre-made parts. The modularity of the strategy allows for the addition or removal of various control elements as a function of the splicing reaction, in the process permitting the cellular location and/or activity state of starting materials and products to be differentiated. The approach is applied to a diverse set of proteins, including the kinase oncofusions BCR/ABL and DNAJB1/PRKACA where dynamic cellular phosphorylation events are dissected, revealing distinct phases of signaling and identifying molecular players connecting the oncofusion to cancer transformation as novel therapeutic targets of cancer cells. We envision that the tools and control strategies developed herein will allow the activity of both naturally occurring and designer proteins to be harnessed for basic and applied research.

## Introduction

Alterations in cellular phenotypes are driven by dynamic changes in the activity of proteins involved in signal transduction and gene regulation processes^1, 2^. Understanding how such changes propagate through associated molecular pathways requires tools that operate on relevant timescales, which can often be in the minutes to hours range^3, 4^. Inducible gene expression approaches such as Cre-Lox and Tet-On/Off systems may not have the temporal resolution to address changes that occur quickly^5, 6^, and while small molecule probes can allow for extremely rapid and specific perturbation (typically inhibition) of a system^3, 7^, the availability of such tool compounds is limited by the druggability of specific targets^8^. This issue of generalizability extends to strategies based on the incorporation of unnatural photocaged amino acids^9, 10^ which, while offering exquisite temporal control, require that the protein in question have the requisite structure-activity characteristics needed for such an approach^11^. Thus, there remains a need for broadly applicable control tools that allow seamless post-translational manipulation of protein structure and function, and that are flexible with respect to the type of control elements one might want to deploy in each context.

In principle, conditional protein trans-splicing (CPS) systems offer an attractive route to the post-translational manipulation of protein function^12–17^. These technologies harness split inteins, which upon complementation, support the ligation of flanking polypeptides (‘exteins’) and in the process remove themselves from the final splice product (Extended Data Fig. 1a). Recent studies into the way intein fragments associate and fold have led to the design of a CPS approach based on the use of genetically ‘caged’ split inteins^18, 19^. In this system, partial sequences from the complementary intein fragments are appended via flexible linkers such that the native N- and C-terminal split inteins (IntN and IntC, respectively) become reversibly trapped in an intramolecular folded state that blocks complementation. Protein trans-splicing is induced upon removal of the extensions by proteolysis^18^, or by bringing the caged split inteins into proximity^19^. The latter scenario involves a domain swapping rearrangement within the heterodimeric complex, a process that triggers protein splicing (Extended Data Fig. 1b). Critically, the final splice product carries no extraneous protein domains associated with the splicing apparatus, a feature that can, in principle, be leveraged to install transient functional control elements within the starting materials.

**Fig. 1.**
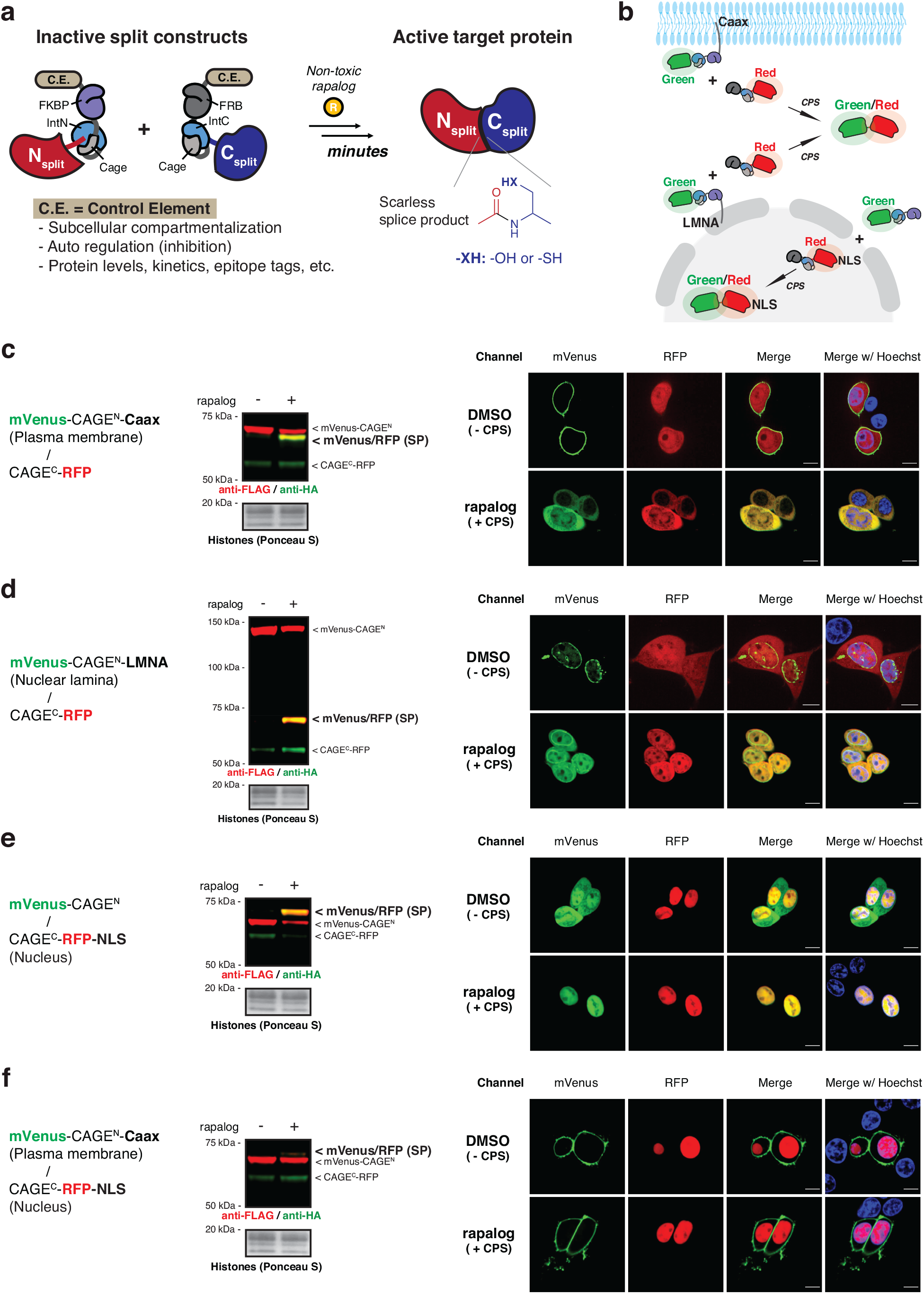
CAGE modules regulate the subcellular location of splice products via control elements. **a,** Schematic of the post-translational activation of a target protein via proximity-triggered CPS using CAGE modules that tightly control the inactive split constructs by appending control elements. **b,** Schematic of the subcellular translocation of mVenus (green) after proximity-triggered CPS with CAGE^C^-RFP (red). The control elements fused to the CAGE modules or exteins regulate the subcellular locations of the starting constructs and the splice product. **c, d, e, f,** Left – immunoblots of the mVenus/RFP splice product (SP) in co-transfected HEK293T cells used for live cell imaging (right). Right – live cell fluorescence imaging of mVenus and RFP localization in HEK293T cells co-expressing mVenus-CAGE^N^ and CAGE^C^-RFP, treated with either DMSO or rapalog (100 nM) for 10 hours. mVenus (green) and RFP (red) were visualized by their intrinsic fluorescence, and Hoechst 33342 (blue) was used as a nuclear marker. Scale bars, 10 µm. ‘Caax’ in (**c**), (**f**) and ‘LMNA’ in (**d**) were fused to the C-terminus of mVenus-CAGE^N^, and ‘NLS’ in (**e**), (**f**) was fused to the C-terminus of CAGE^C^-RFP. Data in (**c**-**f**) (left) represent *n*=3 independent experiments and data in (**c**-**f**) (right) is representative of *n*=12 independent measurements. CAGE^N^ refers to ‘NrdJ-1N^Cage^-FKBP’, and CAGE^C^ refers to ‘FRB-NrdJ-1C^Cage^’ in (**c**-**f**).

In the present study, we incorporate caged split inteins and various control elements into devices that allow post-translational assembly of cellular proteins (Fig. 1a). We show that this strategy permits both the cellular localization and catalytic activity of target proteins to be altered in response to a small molecule trigger. In the case of oncogenic fusion proteins, we demonstrate that this allows dynamic cellular phosphorylation events to be observed, revealing distinct phases of signaling.

## Results

### CAGE modules regulate splice products by altering subcellular localization

Two different caged split inteins were integrated into our system, each responsive to a non-toxic version of rapamycin (rapalog) via an induced proximity mechanism. In addition, to the previously described conditional version of the *Npu* split intein^19^ (Extended Data Fig. 1c, d), which requires a cysteine residue at the splice junction to function, we also incorporated an inducible version of the *NrdJ-1* split intein into the reaction manifold. The latter requires a serine at the splice junction and was thus expected to increase the range of targets amenable to the approach. A rapalog-inducible CPS system based on the split *NrdJ-1* intein was developed following structure-guided optimization of the appended caging sequences on the IntN and IntC fragments (Extended Data Fig. 2a-e). Hereafter, we refer to the caged split inteins and associated FKBP and FRB heterodimerization domains as CAGE^N^ and CAGE^C^. In addition to allowing rapalog-dependent induction of protein trans-splicing in mammalian cells, the *NrdJ-1* cage system was also promiscuous with respect to extein sequences around the splice junction (Extended Data Fig. 2f, g), a property expected to expand the number of potential insertion sites of the CAGE system in target proteins.

**Fig. 2.**
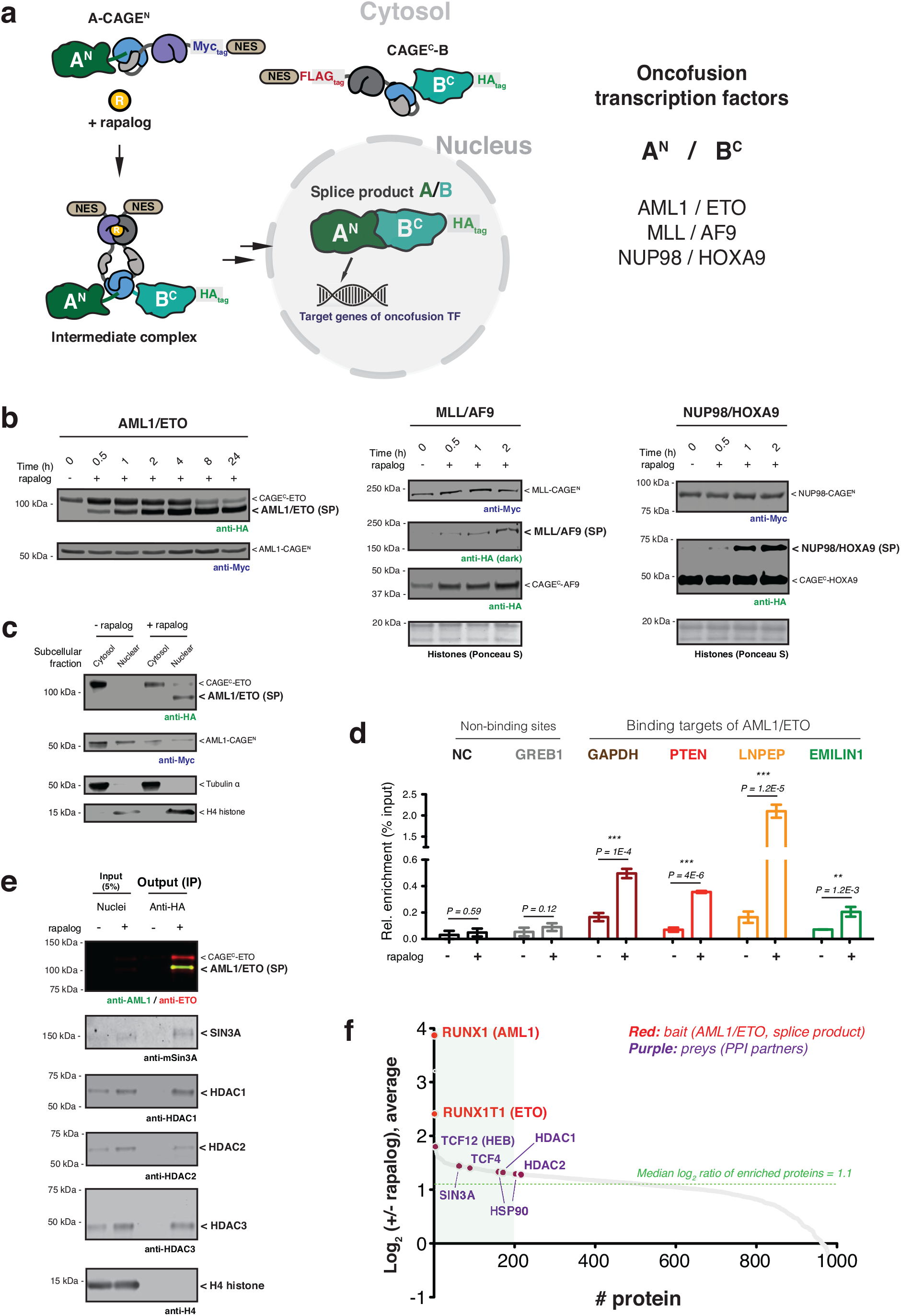
Conditional generation of oncofusion transcription factors. **a,** Schematic showing use of control elements to achieve differential subcellular compartmentalization of A-CAGE^N^ and CAGE^C^-B constructs and the A/B splice product. Oncofusion transcription factors AML1/ETO, MLL/AF9, and NUP98/HOXA9 are generated by proximity-triggered CPS with this strategy. **b,** Immunoblots of splice products (SP, HA tagged) generated in HEK293T cells co-expressing CAGE^N^ (Myc-tag) and CAGE^C^ (HA-tag) constructs. Cells were treated with DMSO or rapalog (100 nM) for indicated time-points. Left – AML1/ETO. Middle – MLL/AF9. Right – NUP98/HOXA9. **c,** Immunoblots of the HA-tagged AML1/ETO splice product generated in HEK293T cells co-expressing AML1-CAGE^N^ (Myc-tag) and CAGE^C^-ETO (HA-tag) treated with rapalog (100 nM) for 18 hours followed by a subcellular fraction to separate cytosolic and nuclear proteins. Tubulin and histone H4 serve as cytosolic and nuclear markers, respectively. **d,** Genomic binding of HA-tagged AML1/ETO splice product. Stable HEK293T cells expressing AML1-CAGE^N^ and CAGE^C^-ETO constructs were treated with DMSO or rapalog (100 nM) for 18 hours. ChIP-qPCR was then used to amplify chromatin derived from immunoprecipitations with anti-HA antibody (*n*=6, mean and s.e.m.). Statistical significance was determined by two-way student’s t-tests: *P < 0.05; **P < 0.005; ***P < 0.001. **e,** Protein binding to the AML1/ETO splice product determined by anti-HA co-immunoprecipitation. Stable HEK293T cells expressing AML1-CAGE^N^ and CAGE^C^-ETO constructs were treated with DMSO or rapalog (100 nM) for 18 hours. Nuclei were isolated and an anti-HA co-immunoprecipitation performed. Proteins known to interact with the native AML1/ETO fusion were probed using the indicted antibodies. H4 serves as a negative control. **f,** Quantitative proteomics analysis of co-enriched proteins following anti-HA immunoprecipitation from nuclei isolated from stable HEK293T cells expressing AML1-CAGE^N^ and CAGE^C^-ETO constructs and treated with DMSO or rapalog. The green box in the waterfall plot indicates the top 20% of enrichment following the induction of PTS. All data points represent the mean ratios (*n*=2, biological replicate runs). Data in (**b**), (**c**), and (**e**) are representative of *n*=3 independent experiments. In panels (**b**-**f**), CAGE^N^ refers to ‘NrdJ-1N^Cage^-FKBP-NES’, and CAGE^C^ refers to ‘NES-FRB-NrdJ-1C^Cage^’.

With two complementary CAGE systems in hand, we next asked whether the intrinsic bond making, and bond breaking features of protein trans-splicing could be harnessed to manipulate cellular protein localization. To test this, model fluorescent proteins mVenus and RFP were fused to CAGE^N^ and CAGE^C^, respectively, along with a series of cellular localization elements whose position within the constructs was expected to result in a change in cellular localization of mVenus upon splicing (Fig. 1b). Consistent with our designs, rapalog treatment of HEK293T cells co-expressing cognate pairs of CAGE constructs led to a splicing-dependent redistribution of mVenus signal from either the plasma membrane, nuclear envelope, or cytosol (Fig. 1c-e and Supplementary Fig. 1). Notably, protein trans-splicing did not occur when the two CAGE constructs were compartmentalized in the cell such that rapalog-induced proximity was prevented (Fig. 1f and Supplementary Fig. 1).

Encouraged by the results with the model fluorescent proteins, we next applied the CAGE system to control the assembly and cellular localization of more challenging protein targets, namely the oncofusion transcription factors, AML1/ETO^20^, MLL/AF9^21^ and NUP98/HOXA9^22^. These fusion proteins result from chromosomal translocations and are associated with various types of leukemia^23, 24^. In each case, CAGE constructs were designed to allow the assembly and nuclear localization of the proteins from pre-made fragments in response to rapalog treatment (Fig. 2a). Notably, use of the *NrdJ-1* cage system permitted the native amino acid sequences at the fusion junctions to be used in the traceless assembly of the oncofusion proteins, whereas the *Npu* cage system required the incorporation of a 2-3 amino acid ‘scar’ to facilitate the slicing reaction (Extended Data Fig. 3a-e and Extended Data Fig. 4a, g). Gratifyingly, we observed dose and time-dependent generation of all three oncofusions upon rapalog treatment of cells expressing the relevant *NrdJ-1* and *Npu* CAGE pairs (Fig. 2b, Extended Data Fig. 3f-h, and Extended Data Fig. 4b-d, h-j). Moreover, as per our designs, we observed nuclear localization of all three oncofusion proteins due to removal of the nuclear export sequences present within the starting materials (Fig. 2c, Extended Data Fig. 3i, and Extended Data Fig. 4e, f, k, l). In the case of the AML1/ETO, a series of genomic and proteomics studies were performed to confirm that the splice product binds to known genomic targets of the oncofusion^25–27^ (Fig. 2d and Extended Data Fig. 5a), and associates with known nuclear protein partners linked to transcriptional regulation^25, 28, 29^ (Fig. 2e, f and Extended Data Fig. 5b-d). Collectively, these data indicate that the CAGE system can be used to control the assembly and cellular localization of target proteins including large oncofusion proteins associated with dysregulation of gene expression in cancer.

**Fig. 3.**
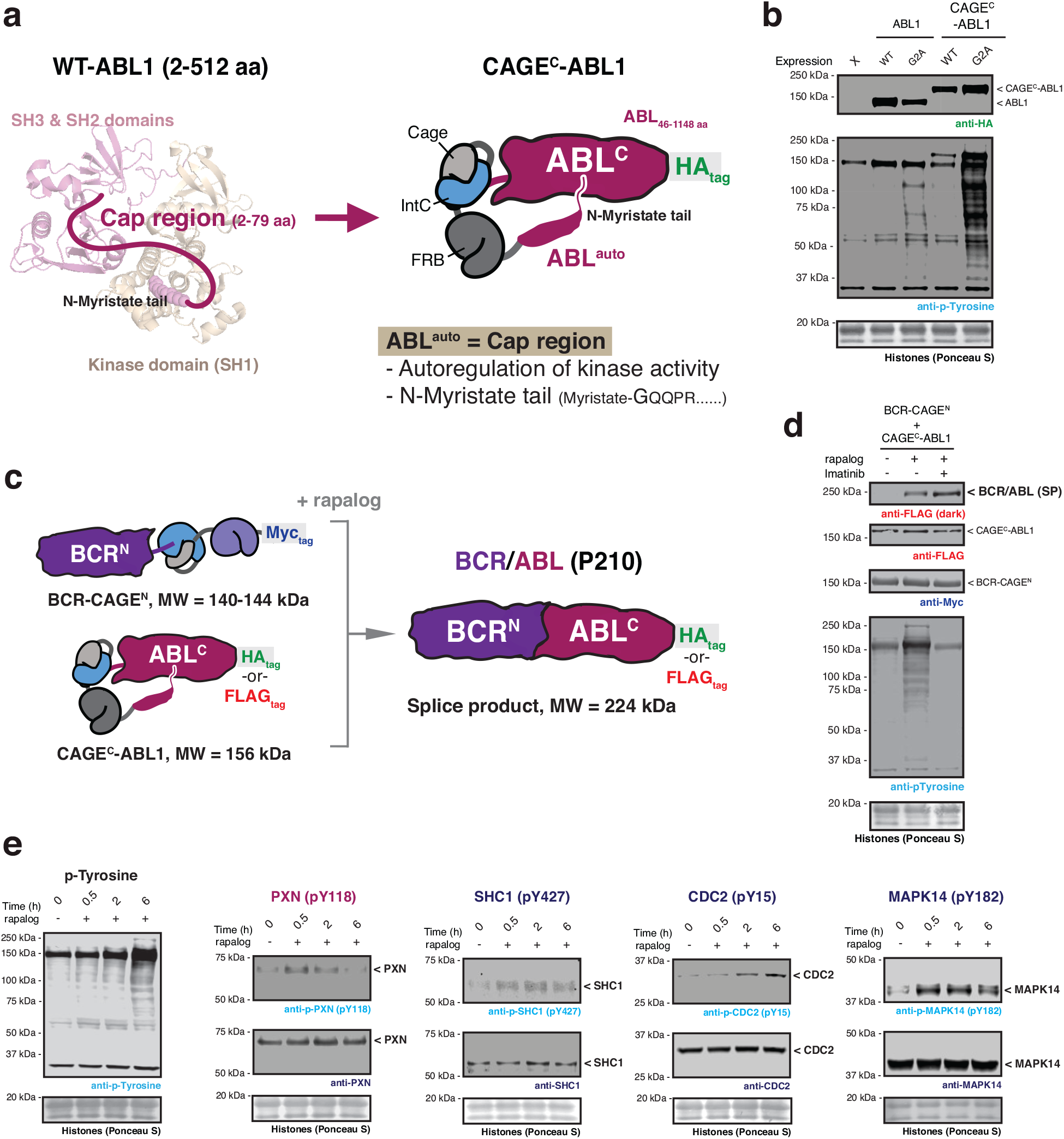
Post-translational generation of BCR/ABL allows dissection of dynamic cellular phosphorylation events. **a,** Left – crystal structure of ABL1 kinase (PDB: 2FO0). The unstructured cap region which connects the SH3 domain to the N-terminal myristoyl group is shown as a cartoon. The myristoyl modification binds to an allosteric site in the C-lobe of the kinase domain. Right – schematic depicting the design of the autoinhibited CAGE^C^-ABL1 construct. **b,** Immunoblots of the basal tyrosine phosphorylation levels in HEK293T cells expressing wild-type or G2A mutant of ABL1 and CAGE^C^-ABL1 constructs. **c,** Schematic of proximity-triggered CPS between complementary BCR-CAGE^N^ and CAGE^C^-ABL1 constructs to generate the BCR/ABL oncofusion. Molecular weights of the splice product and CAGE constructs are indicated. The CAGE^C^-ABL1 construct is kept in an inactive state prior to splicing through use of a *N*-myristoyl control element. **d,** Immunoblots of BCR/ABL (FLAG-tag) splice product (SP) and global tyrosine phosphorylation levels in HEK293T cells co-expressing BCR-CAGE^N^ (Myc-tag) and CAGE^C^-ABL1 (FLAG-tag). Cells were treated with DMSO or rapalog (100 nM) or rapalog and Imatinib (10 µM) for 18 hours prior to analysis. **e,** Immunoblots of the tyrosine phosphorylation levels in HEK293T cells co-expressing BCR-CAGE^N^ and CAGE^C^-ABL1. Cells were treated with rapalog (100 nM) for indicated time-points before analysis with the indicated antibodies. Data in (**b**), (**d**), and (**e**) are representative of *n*=3 independent experiments. CAGE^N^ refers to ‘NpuN^Cage^-FKBP,’ and CAGE^C^ refers to ‘ABL^auto^-FRB-NpuC^Cage^’.

**Fig. 4.**
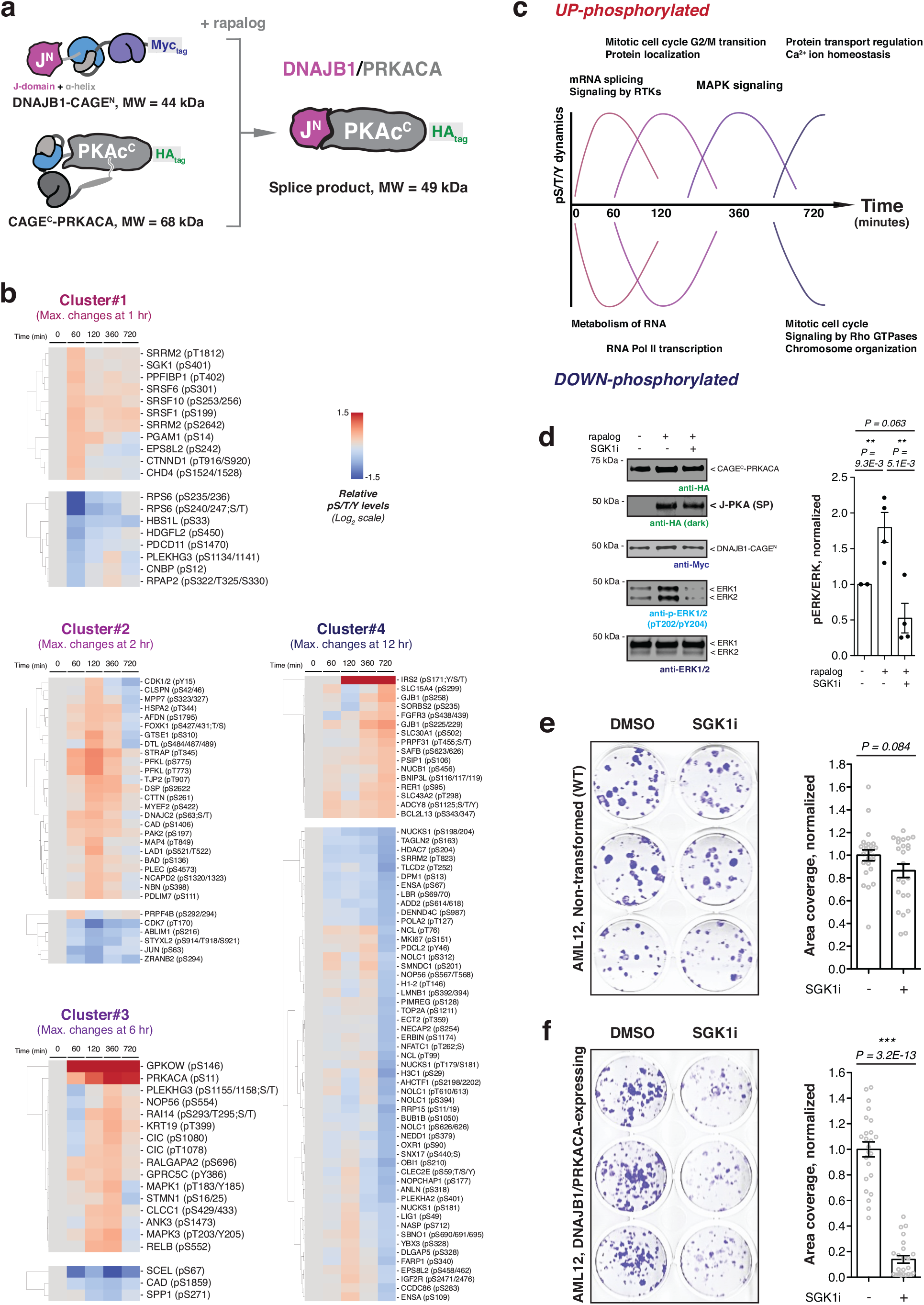
Dynamic cellular phosphorylation events induced by post-translational generation of DNAJB1/PRKACA. **a,** Schematic showing DNAJB1/PRKACA generated by proximity-triggered CPS between complementary DNAJB1-CAGE^N^ and CAGE^C^-PRKACA constructs. Molecular weights of the splice product and the CAGE constructs are indicated, and the *N*-myristate tail in CAGE^C^-PRKACA is proposed to be involved in autoregulation. **b,** Quantitative phospho-proteomics analysis as function of time following the addition of rapalog (100 nM) to AML12 cells co-expressing DNAJB1-CAGE^N^ and CAGE^C^-PRKACA. Phosphorylation sites in the proteomics dataset are grouped into four clusters based on the time taken to reach the average maximum change. Each cluster was subcategorized into ‘UP- and DOWN-phosphorylated’ sites on identified proteins. Data were generated from *n*=3 independent biological replicates. **c,** Summary of the dynamic protein phosphorylation phases downstream of DNAJB1/PRKACA activation as revealed using the CAGE system. **d,** Left – DNAJB1/PRKACA splice product (SP) generation in AML12 cells co-expressing DNAJB1-CAGE^N^ and CAGE^C^-PRKACA. Cells were treated with DMSO, rapalog (100 nM), or rapalog with SGK1 inhibitor (GS-9007, SGK1i, 5 µM) for 6 hours, followed by immunoblotting using the indicated antibodies. Right – the bar charts showing quantified pERK levels of indicated treatments normalized by the ERK intensity (*n*=4, means and s.e.m). **e, f,** Left – representative wells for non-transformed (**e**) and DNAJB1/PRKACA-expressing (**f**) AML12 cells treated with DMSO or GS-9007 (5 µM) for 10 days and stained with crystal violet. Right – clonogenic cell growth of non-transformed (**e**) and DNAJB1/PRKACA-expressing (**f**) AML12 cells quantified by the area covered by colonies and normalized by the average colony area of cells in the DMSO treatment of each plate (*n*=24, means and s.e.m). CAGE^N^ refers to ‘NrdJ-1N^Cage^-FKBP,’ and CAGE^C^ refers to ‘PKAc^auto^-FRB-NrdJ-1C^Cage^’. In panels (**d**-**f**), statistical significance was determined by two-way student’s t-tests: *P < 0.05; **P < 0.005; ***P < 0.001.

### Autoregulated CAGE modules activate BCR/ABL upon protein trans-splicing

Next, we asked whether our approach could be used to control the activity of an enzyme. To explore this, we turned to the oncofusion protein, BCR/ABL, a constitutively active protein tyrosine kinase that drives chronic myeloid leukemia (CML) and that has served as a paradigm for the development of targeted cancer therapy^30^. This oncofusion results from a translocation between chromosomes 9 and 21, also known as the Philadelphia chromosome^31^, resulting in part of the breakpoint cluster region (BCR) protein being linked to the Abelson 1 (ABL1) protein tyrosine kinase. Critically, the fusion results in the loss of a key N-terminal autoregulatory region within ABL1, resulting in de-repression of its kinase activity and, ultimately, mitogenesis^32^. The nature of BCR/ABL activation presented an interesting challenge for extending the scope of our CAGE system since simply splitting the oncoprotein at the fusion junction (as was done in the previous examples) would not eliminate aberrant kinase activity. As a potential solution to this problem, we conceived an intramolecular, allosteric control strategy in which the CAGE apparatus is effectively nested between the ABL1 autoregulatory region and the remainder of the protein (Fig. 3a, c and Extended Data Fig. 6a). We anticipated that this would keep the kinase in the “off-state” until the splicing reaction, which would simultaneously remove this autoregulatory control element whilst ligating the BCR region. To test this idea, we generated a series of constructs of the general form ABL^auto^-linker-CAGE^C^-ABL^C^, where the length of the autoregulatory region (ABL^auto^) and the adjacent linker were varied (Extended Data Fig. 6b). Screening of these candidates led to the identification of an optimal arrangement that exhibited comparable levels of basal kinase activity, as judged by global phosphotyrosine (pTyr) levels, to the wild-type control protein (Extended Data Fig. 6c, d). Further examination revealed that the ABL^auto^ region in the construct was subject to *N-* myristoylation (Extended Data Fig. 6f, g), which is known to be critical for allosteric maintenance of basal ABL kinase activity^33^. Importantly, autoregulation of ABL kinase activity was lost in a mutant version of the CAGE construct, unable to accept the myristoyl modification (Fig. 3b).

We next asked whether the generation of BCR/ABL by proximity-induced CPS would lead to the activation of protein tyrosine kinase activity. Gratifyingly, upon co-expression of the complementary BCR- and ABL1 CAGE constructs, we observed rapalog-dependent generation of the BCR/ABL splice product accompanied by a clear increase in global pTyr levels (Fig. 3d and Extended Data Fig. 6e, h, i). Importantly, this increase was not observed in the presence of the ABL inhibitor Imatinib (Fig. 3d). From this, we conclude that the observed increase in global phosphorylation is due to the generation of BCR/ABL via proximity-triggered CPS. To explore cellular pTyr dynamics upon acquisition of onco-kinase activity, we carried out time-resolved quantitative phospho-proteomics (Supplementary Fig. 2a, b). This analysis revealed 39 sites in 37 proteins whose pTyr levels reproducibly increased over 6 hours in a manner that satisfied our stringency criteria (Extended Data Fig. 7a, b and Supplementary Fig. 2c). This list included multiple pTyr sites within BCR/ ABL itself, as well as pTyr sites within several receptor protein tyrosine kinases and cyclin dependent kinases. Broadly speaking, the identified pTyr sites could be divided into those that reach a maximum within 2 hours of initiation, versus those that accumulate more slowly (Extended Data Fig. 7b). Among the former group we were surprised to see that several proteins, such as the cytoskeletal signaling protein paxillin (pY118), exhibited an acute increase in phosphorylation which then decreased at later timepoints. This differential dynamic behavior was verified for representative examples using western blotting analysis (Fig. 3e). Notably, use of a Tet-On system for inducible BCR/ABL expression in HEK293T cells did not lead to comparable changes in pTyr dynamics at the earlier timepoints following doxycycline treatment (Extended Data Fig. 7c).

### Time-resolved DNABJ1/PRKACA activation deconvolutes acute changes in cellular signaling

Given the success with BCR/ABL, we were keen to determine whether our CAGE system could be applied to another oncofusion involving a protein kinase. For this, we chose the DNAJB1/PRKACA fusion protein which is the key driver in fibrolamellar hepatocellular carcinoma (FL-HCC), a rare cancer of the liver^34, 35^. This oncofusion results from a deletion in chromosome 19, resulting in the in-frame fusion of the J domain of heat shock protein DNAJB1 and most of the catalytic subunit of cAMP-dependent protein kinase (PRKACA)^34^. Reminiscent of BCR/ABL, the genetic fusion results in loss of the first 14 residues from PRKACA which, notably, includes an autoregulatory *N-*myristoylation modification^36^ (Extended Data Fig. 8a, b). While DNAJB1/PRKACA is regarded as the central player in FL-HCC oncogenesis, many questions remain regarding how cellular signaling is altered by this chimeric protein during cell transformation^37–40^. Consequently, we imagined that the ability to control the synthesis of the chimera using our CAGE system could provide new insights into these molecular mechanisms.

Informed by the crystal structure of DNAJB1/PRKACA^41, 42^, we designed a *NrdJ-1* CAGE system in which the splicing junction was located within the unstructured linker region immediately following the N-terminal alpha-helix of PRKACA (Fig. 4a and Extended Data Fig. 8b-d). A similar regulation strategy was adopted to that successfully employed in the BCR/ABL system. Thus, the CAGE apparatus was nested between the autoregulatory region (PKAc^auto^) and PRKACA (Extended Data Fig. 8c, d). Metabolic labeling studies confirmed that the PKAc^auto^ control element in the CAGE^C^-PRKACA construct was *N*-myristoylated (Extended Data Fig. 8e, f). Immunoblotting using an antibody against phospho-PKA substrates revealed that HEK293T cells co-expressing the complementary DNAJB1- and PRKACA CAGE constructs did not have increased levels of phosphorylation compared to control cells lacking the CPS apparatus (Extended Data Fig. 8g). By contrast, rapalog-induced generation of the DNAJB1/PRKACA splice product resulted in a change in the global PKA phosphorylation signature similar to that of positive control cells expressing the oncofusion kinase (Extended Data Fig. 8g, h).

Next, we imported the DNAJB1/PRKACA CAGE system into AML12 cells, non-transformed murine hepatocytes that have previously been used for studying this oncofusion protein^37, 38^. As anticipated, the DNAJB1/PRKACA product was generated in a time and dose-dependent manner upon rapalog treatment of a stable AML12 cell line co-expressing the CAGE constructs (Extended Data Fig. 8i, j). Co-immunoprecipitation studies confirmed that the DNAJB1/PRKACA splice product associates with the known interaction partner, chaperonin heat shock protein 70 (HSP70)^37^ (Extended Data Fig. 8k). We then carried out time-dependent quantitative phosphoproteomics analyses in both HEK293T and AML12 cells following induction of protein splicing. Analysis of the HEK293T dataset revealed 245 phosphopeptides from 230 proteins whose phosphorylation levels reproducibly changed over 12 hours, as exemplified by the phosphorylation at PRKACA and HSP70 (Supplementary Fig. 2c and Supplementary Fig. 3a, b). Analysis of the dataset using the Kinase Library platform^43^ revealed that the basophilic motif and the AGC (PKA, PKG and PKC) kinase family target sites are enriched in the UP-phosphorylated group, whereas the DOWN-phosphorylated group align with proline-directed motif sites (Supplementary Fig. 3c, d). Broadly similar results were obtained from the AML12 cell line, in this case 138 phosphopeptides from 127 proteins reproducibly changed during the time course (Supplementary Fig. 4a-d).

To further explore the signaling events downstream of DNAJB1/PRKACA generation, we grouped the proteins in the datasets based on phosphorylation dynamics (Fig. 4b and Extended Data Fig. 9). All clusters were characterized by maximal changes of phosphorylation levels, either increased or decreased, at the specific time points after triggering splicing; 1 hour (cluster 1), 2 hours (cluster 2), 6 hours (cluster 3), and 12 hours (cluster 4). GO analysis indicated distinct biological roles for protein members in each cluster (Supplementary Fig. 5a, b). In the mouse liver cells, proteins in cluster 1 are linked to mRNA splicing/receptor tyrosine kinases-associated signaling (phosphorylated) or RNA metabolism (dephosphorylated), cluster 2 associates with mitotic cell transition/protein localization (phosphorylated) or RNA Pol II transcription (dephosphorylated), cluster 3 is linked to MAPK signaling (phosphorylated), and the top terms in cluster 4 are regulation of protein/calcium ion transport (phosphorylated) or mitotic cell cycle/Rho GTPases signaling/chromosome organization (dephosphorylated), respectively (Fig. 4c). Notably, the dynamic phosphorylation events downstream of DNAJB1/PRKACA appear to be context-dependent – the GO terms associated with each cluster are not the same between the HEK293T and AML12 cells two cell types, a divergence highlighted by principal component analysis (PCA) of the phosphoproteomics datasets (Supplementary Fig. 6).

Finally, we wondered if our inducible DNAJB1/PRKACA system could provide insights into the molecular player(s) involved in up-regulation of the MAPK/ERK signaling pathway, which is a hallmark of FL-HCC^37, 38^. With this in mind, we were particularly intrigued by the finding that phosphorylation levels of Serum and glucocorticoid-regulated kinase 1 (SGK1) increased at early time points upon induction of DNAJB1/PRKACA in AML12 cells (Fig. 4b). SGK1, a member of the AGC kinase family, is overexpressed in many cancers where it affects cellular signaling pathways, including MAPK/ERK activation^44–46^. Interestingly, phosphorylation at serine 401 on SGK1 identified in our phospho-proteomic analysis was previously found to be essential for full SGK1 activity^47^, suggesting this protein as a candidate for linking DNAJB1/PRKACA to activation of the MAPK/ERK pathway. Consistent with this idea, elevated ERK phosphorylation was not observed in DNAJB1/PRKACA-activated AML12 cells treated with the SGK1 inhibitor, GS-9007^48, 49^ (Fig. 4d). Furthermore, compared to the non-transformed AML12 cells, the growth of AML12 cells stably expressing DNAJB1/PRKACA and their ERK phosphorylation were selectively inhibited by GS-9007 (Fig. 4e, f and Extended Data Fig. 10a-c). Together, these results implicate SGK1 in connecting the activity of the DNAJB1/PRKACA oncofusion with elevated MAPK signaling in this early FL-HCC model (Extended Data Fig. 10d).

## Discussion

In this study, we have developed strategies for controlling protein function at that post-translational level using caged split inteins. The platform has been successfully applied to multiple targets, including five different native oncofusions of differing sizes and functions, illustrating the generality of the method. The temporal control afforded by our strategy allowed us to follow changes in cellular phosphorylation following the generation of the kinase oncofusions. In the case of BCR/ABL, different phases of pTyr dynamics were uncovered. Perhaps most remarkable was the observation of transient phosphorylation events that occur in the first few hours after initiating splicing, including in receptor tyrosine kinases (RTKs). The fact that we also observed delayed tyrosine phosphorylation of known substrates of these RTKs, including various nuclear ribonucleoproteins, illustrates how our approach can provide insights into the cascade of signaling events emanating from the onset of BCR/ABL kinase activity.

Application of the CAGE system to DNAJB1/PRKACA has provided insights into the signaling pathways perturbed by this oncofusion. Our phospho-proteomics studies indicate that many basophilic motif sites and AGC kinase targets, including serine 11 in WT-PRKACA, undergo phosphorylation following DNAJB1/PRKACA generation. However, it is worth noting that early transient phosphorylation targets were not solely dominated by this single motif and kinase targets. This suggests that DNAJB1/PRKACA may contribute to oncogenesis not only through its kinase activity but also by deregulating signaling landscapes through other mechanisms^37, 39^. Secondly, our research has revealed distinct temporal phases of phosphorylation following DNAJB1/PRKACA generation, including MAPK signaling, which provides insight into the underlying molecular connections between the oncofusion and cancer transformation. With respect to this, we provide evidence that the kinase, SGK1, acts as upstream intermediate connecting DNAJB1/PRKACA activity to the MAPK pathway. Lastly, our research revealed multiple dephosphorylation events after DNAJB1/PRKACA generation in the HEK293T and AML12 cell lines. Notably, we observed the early transient phosphorylation of several phosphatase regulatory subunits after DNAJB1/PRKACA generation, suggesting potential mechanistic links that could be explored in future studies.

The CAGE system described herein possesses several key attributes that can be exploited for controlling protein function. Firstly, since it works at the post-translational level, target fusion proteins can be generated in a dose and time-dependent manner from premade parts. As demonstrated above, this provides opportunities for studying acute changes in cellular physiology upon protein gain of function. Secondly, the method is traceless, meaning that the entire CPS apparatus, as well as any other appended control elements, is removed from the final splice product. This distinguishes the approach from more traditional induced proximity strategies based on chemical or optogenetic inputs where the assembly apparatus is retained in the final complex^50, 51^. Lastly, the fact that amide bonds are both broken and formed during protein trans-splicing provides opportunities for building additional control into the system. In this study, we demonstrated this through the transient introduction of multiple subcellular localization factors and autoregulatory domains with the CAGE constructs. However, we envision that this approach can be extended to other control elements, such as protein domains that bind to other proteins or nucleic acids^52, 53^. More generally, the system provides a platform for the ‘metathesis’ of protein molecules; splicing results in the simultaneous exchange of multiple functional domains through both covalent and non-covalent interactions (Supplementary Fig. 7). This capability should engender the approach to a wide range of synthetic biology applications.

## Acknowledgements

We thank Saw Kyin and Henry H. Shwe at the Princeton University Proteomics and Mass Spectrometry Core Facility. We thank Christina J. DeCoste and Katherine Rittenbach at the Princeton University Flow Cytometry Resource Facility for cell sorting services funded by Rutgers Cancer Institute of New Jersey P30CA072720-5921. We thank Gary S. Laevsky and Sha Wang at the Princeton University Confocal Imaging Facility. We also thank Jennifer M. Miller at the Princeton University Genomics Core Facility. Research reported in this publication was supported by NIH grant R01GM086868 (T.W.M).

## Author contributions

G.L. and T.W.M. conceived the work. G.L. and T.W.M. designed and performed the experiments.

G.L. and T.W.M. wrote and reviewed the manuscript.

## Data and materials availability

All data are available upon request.

**Extended Data** is available for this paper.

Extended Data Fig. 1-10

**Supplementary Information** is available in the online version of this paper.

Supplementary Fig. 1-7

Supplementary Table 1-11

## Methods

### Materials

Alkynyl myristic acid (YnMyr, CAS No. 82909-47-5) was obtained from Click Chemistry Tools. Azo biotin-azide (CAS No. 1339202-33-3) and Imatinib (Cas No.152459-95-5) were obtained from Sigma Aldrich, rapamycin (Cas No.53123-88-9) from ApexBio (Huston, TX), AP21967 (rapalog) from Takara, Doxycycline (Cas No.24390-14-5) from Stem Cell Technologies, and SGK1 inhibitor GS-9007 (Cas No.1426214-51-8) was obtained from Cayman Chemical. All commercially obtained chemicals were dissolved in DMSO and used without further purification.

### Cell culture

HEK293T and AML12 cell lines were purchased from ATCC. HEK293T and stably expressed HEK293T cells were propagated in DMEM (Thermo Fisher) supplemented with 10% v/v fetal bovine serum (FBS, Atlanta Biologicals), 2 mM L-glutamine (Life Technologies), and 1% v/v penicillin-streptomycin (5,000 U/mL, Life Technologies). AML12 and stably expressed AML12 cells were propagated in DMEM/F-12 (Gibco) supplemented with 10% v/v FBS, 0.1% v/v gentamicin (50 mg/mL, Sigma Aldrich), 1% v/v ITS-G (100x, 1.0 mg/mL recombinant human insulin, 0.55 mg/mL human transferrin, 0.67 μg/mL sodium selenite, Gibco), and 0.04 μg/mL dexamethasone (Sigma Aldrich). AML12 and stable AML12 cells were switched to serum-free media for 24 hours before the experiments and treated with the rapalog under the same condition^37^. All cell lines were grown at 37 <C in a 5% CO_2_ humid incubator.

### Transfection

Each plate (6-well or 10 cm, Fisher Scientific) of HEK293T cells at ∼70% confluency was transfected with plasmids encoding ‘POI^N^-CAGE^N^’ and ‘CAGE^C^-POI^C^’ with Lipofectamine 2000 (Thermo Fischer) according to the manufacturer’s protocol - ‘POI’ = Protein of Interest. After 6 hours, the media was aspirated and replaced with fresh media. Transfection was performed for 24 or 48 hours in an incubator at 37 <C in 5% CO_2_.

### Lentiviral production and generation of stable cell lines

Lenti-X 293T cells (Takara) were seeded in 6-well plates (Fisher Scientific) at 1.0 x 10^6^ cells per well and transfected after 24 hours with transfer plasmid (POI^N^-CAGE^N^_IRES2_EGFP or CAGE^C^-POI^C^_IRES2_mCherry in pEF1α backbone, 4 μg) and packaging vectors (pMD2.G 1.2 μg and psPAX 3.6 μg) using Lipofectamine 2000. Following overnight transfection, media was exchanged and allowed to incubate for an additional 24 hours. The viral collection was performed at 24, 48, and 72 hours. Viral media was filtered with a 0.45 μm PES syringe filter (Membrane Solutions), and Polybrene (Sigma #H9268) was added to a concentration of 8 μg/mL before infection of target cell lines.

HEK293T and AML12 cell lines were infected with the 24-hour or 48-hour viral harvests. After 24 hours, cells were allowed to recover by exchanging the media. Serial infections were performed for co-viral transduction of the two CAGE constructs; cells were initially selected with Blasticidin S (20 μg/mL final concentration to HEK293T cells and 2 μg/mL to AML12 cells, Gibco) after the first viral infection, then another virus was added to the cells for the second infection. Then, double-positive cells were isolated from the mix via FACS after being cultured with Blasticidin S.

BCR/ABL Tet-On HEK293T cells were generated by transfection with a Piggybac transposon plasmid encoding for BCR/ABL under the control of a doxycycline-inducible promoter and a Piggybac transposase plasmid (a gift from the Kadoch lab, Dana Farber Cancer Institute) (2:1 ratio). Lipofectamine 3000 (Thermo Fischer) was used for the transfection according to the manufacturer’s protocol, transformed cells were selected by being cultured with 10 μg/mL Blasticidin S after 30 hours of transfection.

### Fluorescence-activated cell sorting (FACS)

After co-viral transduction followed by selection with Blasticidin S, cells were collected after trypsin treatment (Trypsin-EDTA 0.05%, Gibco), washed twice with FACS buffer (2% FBS in PBS), and resuspended in FACS buffer at a 1 x 10^6^ cells/mL concentration. Cells were filtered through the cell strainer (40 μm, Fisher Scientific) and sorted using a FACSAria Fusion cell sorter (BD Bioscience) at the Princeton Flow Cytometry Resource Facility. Cells expressing both GFP and mCherry were collected as single clones in 96-well plates or in a common tube and expanded in DMEM media until used for testing CPS modules in cells. The GFP was excited by the 488 nm laser, with emitted fluorescence detected through a 530/30 bandpass filter, and the mCherry was excited by the 561 nm laser, and emission was detected through a 610/20 bandpass filter. The cell sorter was equipped with a 100-micron nozzle, and the sheath pressure was maintained at 20 psi.

### Cloning

All cloning was performed using the Phusion High-Fidelity DNA polymerase (New England Biolabs) and primers from IDT or Sigma Aldrich. Plasmids of the general forms ‘POI^N^-CAGE^N^’ and ‘CAGE^C^-POI^C^’ were cloned into vector backbones using Gibson Assembly (New England Biolabs) following the manufacturer’s instructions. The encoded constructs were controlled by the CMV promoter (strong expression when transiently transfected, pCMV5 vector backbone) and sv40 promoter (weak expression when transfected, pSG5 vector backbone). Point mutations were installed using inverse PCR. For generating lentiviral vectors for constitutive expression of CAGE constructs, the genes incorporating POI^N^-CAGE^N^ with IRES2_EGFP or CAGE^C^-POI^C^ with IRES2_mCherry were cloned into the 3^rd^ generation lentiviral vector via restriction enzyme digestions using ClaI and MluI (New England Biolabs) followed by ligation using T4 DNA ligase (New England Biolabs). Plasmid sequences were verified by Sanger sequencing (Genewiz) using universal primers. Amino acid sequences for the encoded proteins are provided in Supplementary Table 1. In addition, the primer sequences used for point mutations of caging sequence in NrdJ-1 CAGE modules were provided in Supplementary Table 2.

### Live cell imaging

HEK293T cells co-expressing ‘mVenus-CAGE^N^’ and ‘CAGE^C^-RFP’ were seeded onto a 35 mm dish with a 14 mm glass diameter (MatTek) and treated with either rapalog or DMSO for 8-10 hours. Next, the cells were stained with Hoechst 33342 (1:2,000 dilution, Invitrogen) for 5 minutes and washed twice with Live Cell Imaging Solution (Thermo Fisher). The stained cells were imaged using a Nikon CSU-21 spinning disk confocal microscope with NIS Elements (version 5.2) software. Excitation of mVenus was perfromed using a 488 nm laser, RFP was excited using a 561 nm laser, and Hoechst was excited using a 405 nm laser. Images were captured with a 60x Plan Apo λ Oil objective and a Hamamatsu ORCA-Flash4.0 sCMOS camera.

### Subcellular fractionation

1.0 x 10^7^ HEK293T cells transfected with plasmids encoding ‘POI^N^-CAGE^N^’ and ‘CAGE^C^-POI^C^’ and treated with DMSO or rapalog (100 nM) were lysed by hypotonic lysis in 1 mL RSB buffer (10 mM Tris, 15 mM NaCl, 1.5 mM MgCl_2_, EDTA-free Halt protease inhibitor cocktail, pH 7.6) for 10 minutes on ice. The crude nuclei were isolated by centrifugation at 400 g for 5 minutes at 4 <C. After collecting the supernatant as a cytosolic fraction, the nuclei were resuspended in 1 mL RSB buffer, homogenized with 10 strokes of a loose pestle Dounce homogenizer, and pelleted at 400 g for 5 minutes at 4 <C. The nuclei were resuspended in 1 mL RSB buffer and centrifuged at 700 g for 5 minutes at 4 <C. Finally, the nuclei were resuspended in 300 μL of lysis buffer (Tris 20 mM, NaCl 150 mM, 0.5% v/v SDS, EDTA-free Halt protease inhibitor cocktail, pH 7.6), sonicated (Fisher Scientific FB-505, Amplitude: 25%, No pulse, Time: 15 seconds x 2 times), and insoluble debris cleared by centrifugation at 17,000 g for 10 minutes at 4 <C. Alternatively, the isolated nuclei were resuspended in desired buffers for other experiments, such as ChIP or co-immunoprecipitation assays.

### Metabolic labeling of cells with YnMyr and N-myristoylated protein enrichment

Enrichment of N-myristoylated proteins in cells treated with the analog probe was performed using a protocol adapted from the reference^54^ with minor modifications, as described below. HEK293T cells transfected with the plasmid encoding ‘ABL1-HA’, ‘CAGE^C^-ABL1-HA’, or ‘CAGE^C^-PRKACA-HA’ were incubated with culture media containing YnMyr (20 μM) for 24 hours before cell lysis. After treatment, cells were washed with cold PBS (2 x) and lysed on ice using lysis buffer (Tris 20 mM, NaCl 150 mM, 1% v/v SDS, EDTA-free Halt protease inhibitor cocktail, pH 7.6) with sonication (Fisher Scientific FB-505, Amplitude: 30%, No pulse, Time: 8 seconds x 2 times). Lysates were centrifuged at 17,000 g for 10 minutes to remove insoluble material. The supernatant was collected, and protein concentration was measured using the BCA assay (Thermo Fisher). A click mixture containing azido-azo biotin, Tris(2-carboxyethyl)phosphine (TCEP), Tris(benzyltriazolylmethyl)amine (TBTA), and CuSO4 was prepared by combining the reagents in that order. The click mixture was then added to the diluted lysate and the reaction allowed to proceed for 1 hour at room temperature in the dark. The final reaction conditions were as follows: 1 mg/mL protein, 1% SDS, 50 mM Tris pH 8.0, 0.1 mM Azo biotin-azide (stock solution 5 mM in DMSO), 1 mM TCEP (stock solution 50 mM in ddH_2_O, prepared fresh), 0.2 mM TBTA (stock solution 10 mM in DMSO), 1 mM CuSO_4_ (stock solution 50 mM in ddH_2_O, prepared fresh). The samples were vortex-mixed briefly with the addition of methanol (3 volumes), chloroform (0.75 volume), and ddH_2_O (2 volumes), then centrifuged at 17,000 g for 10 minutes to pellet precipitated proteins. After removing both the top and bottom layers, protein pellets were washed twice with methanol (2.5 volumes) and air-dried. After protein precipitation, the pellet was resuspended in PBS containing 1% SDS supplement with EDTA-free Halt protease inhibitor. Once the pellet was completely dissolved, the samples were diluted with 1% SDS in PBS to 1 mg/mL protein concentration.

Streptavidin agarose resin (Thermo Scientific) was washed with 1% v/v SDS PBS (2 x). Typically, 50 μL of resin slurry was used for 1 mg lysate. The agarose beads were added to the samples, and the affinity enrichment carried out overnight at 4 <C or 2 hours at room temperature. The supernatant was removed, and the beads were washed with 1% SDS in PBS (3 x) followed by PBS (2 x). Enriched proteins were eluted by either heating the beads with 2X Laemmli buffer containing 4% beta-mercaptoethanol (βMe) at 95 <C for 10 minutes or by incubating the beads with 25 mM of Sodium dithionite (Sigma Aldrich) and 1% SDS in PBS at room temperature for 30 minutes (2 x). Samples were then prepared for SDS-PAGE analysis below.

### Immunoblotting

Cells were harvested by scraping, pelleted by centrifugation, washed twice with PBS, and lysed in 8 M urea, 50 mM NH_4_HCO_3_ pH 8.0 or TBS (50 mM Tris, 150 mM NaCl, pH 7.6) containing 1% SDS supplement with EDTA-free Halt protease inhibitor and phosphatase inhibitor cocktails (Thermo Scientific) with sonication (Amplitude: 20%, No pulse, Time: 6 seconds) at 4 <C. Cells were then cleared by centrifugation, and the supernatant was diluted into 4x Laemmli buffer containing 8% beta-mercaptoethanol (βMe) as reducing agents. Samples were prepared for SDS-PAGE by heating to 95 <C for 10 minutes, cooled to room temperature, resolved on 6%, 8%, and 4-12% Bis-Tris protein gels, and transferred onto nitrocellulose membranes by standard immunoblotting methods. Membranes were blocked in 3% BSA in TBS containing 0.1% tween-20 (TBST) and probed with primary antibodies (see Supplementary Table 3). After the overnight incubation of membranes with primary antibodies, the secondary donkey anti-rabbit and donkey anti-mouse (Licor) were used at 1:10,000 dilution in 3% BSA-containing TBST and incubated for 1 hour prior to washing and imaging on a Licor infrared scanner. Whole protein bands were stained via Ponceau S staining solution (Thermo Fisher). Densitometry measurements were performed with ImageJ software.

Time- and dose-dependent rapalog treatment studies or doxycycline treatments (200 ng/mL) were performed in cells 24 hours after transient transfection of plasmids encoding ‘POI^N^-CAGE^N^’ and ‘CAGE^C^-POI^C^’ or stable cells. The indicated concentration of rapalog (stock solution 1 mM in DMSO) or equivalent DMSO in appropriate media was added to cells in 6-well plates or 10 cm dishes. Following the indicated incubation, cells were lysed in 8 M urea, 50 mM NH_4_HCO_3_ pH 8.0, or TBS containing 1% SDS supplement with EDTA-free Halt protease inhibitor and phosphatase inhibitor cocktails (Thermo Scientific) and processed for immunoblotting as indicated above.

### ChIP sample preparation

HEK293T cells stably expressing the genes encoding ‘AML1^N^-CAGE^N^’ and ‘CAGE^C^-ETO^C^’ were treated with DMSO or rapalog (100 nM) followed by fixing the cells in 1% formaldehyde for 8 minutes. After quenching the excess formaldehyde with Glycine (∼ 0.125 M final, from 2.5 M stock), cells were washed with cold PBS (2 x) and lysed by following the subcellular fractionation protocol to isolate the nuclei. Then, the nuclear pellets were resuspended in high salt buffer (50 mM HEPES-NaOH, 500 mM NaCl, 1 mM EDTA, 0.1% Sodium deoxycholate, 1% Triton-X, 0.05% SDS, EDTA-free Halt protease inhibitor, pH 7.5) and sonicated (Amplitude: 25%, Pulse: 15 seconds on / 45 seconds off, Time: 2 minutes – 8 cycles) at 4 <C. The samples were centrifuged at 17,000 g for 10 minutes to remove insoluble material. Each ChIP was performed with 1 million cells, and chromatin samples were incubated overnight at 4 <C with anti-HA magnetic beads (Pierce, 88836) washed with high salt buffer (3 x) before incubation. The immobilized protein-DNA complexes were washed 3 times with high salt buffer, 2 times with low salt buffer (10 mM Tris-Cl, 250 mM LiCl, 1 mM EDTA, 0.5 % NP40, 0.5% sodium deoxycholate, pH 8.0), and 1 time with TE buffer (10 mM Tris-Cl, 1 mM EDTA, pH 8.0). Elution and reverse-crosslinking were carried out in the elution & de-crosslink buffer (50 mM Tris-Cl, 10 mM EDTA, 1% SDS, pH 8.0) supplemented with RNAse A and Proteinase K at 65 <C for 4 hours. Inputs without affinity pulldown with antibodies were also reverse-crosslinked by the same method. The supernatants were collected and purified via PCR purification and gel extract columns (Tomas Scientific), and DNA was eluted from columns with TE buffer.

### ChIP-qPCR and data analysis

ChIP samples and inputs (1%) were used for real-time quantitative polymerase chain reaction (RT-qPCR) to get cycle threshold (Ct) values. Quantification of PCR was performed with PowerUp^TM^ SYBR^TM^ Green master mix (Thermo Fisher) using a ViiA 7 Real-Time PCR system (Thermo Fisher). Ct values from *n*=6 (three measurements from two independent experiments) of each sample were normalized by input Ct values, and relative enrichment (% of input) was calculated with standard deviations. In addition, *P* values were calculated by univariate two-sided t-test with the values of rapalog and DMSO treated conditions. The qPCR primer sequences used in this study are provided in Supplementary Table 4, and Ct values of ChIP-qPCR experiments are provided in Supplementary Table 5.

### Co-immunoprecipitation assay (AML1/ETO)

10 cm plates of HEK293T cells stably expressing the genes encoding ‘AML1^N^-CAGE^N^’ and ‘CAGE^C^-ETO^C^’ were treated with DMSO or rapalog (100 nM) for 18 hours and washed with cold PBS (2 x). After hypotonic lysis, nuclei were isolated by following the subcellular fractionation protocol. Then, isolated nuclei samples were resuspended in 450 μL modified buffer A (20 mM HEPES, 1.5 mM MgCl_2_, 150 mM KCl, 0.1% Triton-X, 0.05% SDS, EDTA-free Halt protease inhibitor cocktail, pH 7.5) and benzonase nuclease (4 μL, Millipore) was then added to each condition for 1-2 hours at room temperature (Benzonase nuclease digestion time varies by enzyme batch and must be determined for each experiment. 3 μL of the digest was taken every 30 minutes, and these aliquots were run on a 1% agarose gel, and the digestion efficiency was visualized with ethidium bromide staining). Once digestion was completed, the samples were spun at 1.3 kg for 5 minutes at 4 <C, and the supernatants were collected (fraction S1). 550 μL of modified buffer A was added to the pellets, and each sample was rotated end-over-end at 4 <C for 1-2 hours. The samples were spun at 13,000g for 5 minutes at 4 <C, and the supernatants were collected (fraction S2). Fraction S1 and S2 are combined in a Lo-bind Eppendorf tube after centrifugation (13,000g for 5 minutes), and the protein concentration of each sample was measured using the BCA assay (Thermo Fisher) and diluted with modified buffer A to 0.8 mg/mL protein concentration. 35 μL of anti-HA magnetic beads (Pierce, 88836) were pre-washed twice with modified buffer B (20 mM HEPES, 1.5 mM MgCl_2_, 150 mM KCl, 0.3% Triton-X, 0.1% SDS, EDTA-free Halt protease inhibitor, pH 7.5) and added to the sample. The HA-IP was performed overnight with end-over-end rotation at 4 <C. The remaining samples were kept as inputs. The beads were washed sequentially with 4x modified buffer B and 1x ddH_2_O for 2 minutes each rotation. 120 μL of 1x Laemmli buffer (no reducing agents) was added, and the beads were heated at 98 <C for 10 minutes. After collecting the eluents and adding reducing agents (DTT, 10 mM final), samples were run on a Bis-Tris gel for immunoblotting with indicated antibodies (see Supplementary Table 3).

### Co-immunoprecipitation assay (DNAJB1/PRKACA)

15 cm plates of AML12 cells stably expressing the genes encoding ‘DNAJB1-CAGE^N^’ and ‘CAGE^C^-PRKACA’ were treated with DMSO or rapalog (100 nM) for 24 hours and washed with cold PBS (2 x). Cells were then lysed in TBS (50 mM Tris, 150 mM NaCl, pH 7.6) containing 1% Triton-X supplement with EDTA-free Halt protease inhibitor cocktails (Thermo Scientific) with sonication (Amplitude: 20%, Pulse: 1 second on / 1 second off, Time: 30 seconds) at 4 <C. Lysates were centrifuged at 17,000 g for 10 minutes to remove insoluble material, and protein concentration was measured by BCA assay. Lysates were then diluted to 2 mg/mL protein concentration and transferred to a Lo-bind Eppendorf tube. Next, 20 μL of anti-HA magnetic beads (Pierce, 88836) were pre-washed twice with the lysis buffer above and added to the sample. The HA-IP was performed overnight with end-over-end rotation at 4 <C. The remained samples were kept as inputs. The beads were washed sequentially with 5 times lysis buffer containing 0.05% SDS and 2 times ddH_2_O for 2 minutes each rotation. Next, 75 μL of S-TRAP elution buffer (TEAB 50 mM, 5% SDS, pH 7.1) was added, and the beads were heated at 45 <C for 10 minutes. After collecting the eluents and diluted into 4x Laemmli buffer containing 8% beta-mercaptoethanol (βMe) as reducing agents, samples were run on a Bis-Tris gel for immunoblotting.

### SILAC cell culture

SILAC labeling was performed by growing cells for at least eight passages in lysine- and arginine-free SILAC medium (DMEM, Invitrogen) supplemented with 10% dialyzed fetal calf serum (Thermo Fisher), 2 mM L-glutamine, and 1% P/S. “Light” and “heavy” media were supplemented with natural lysine and arginine (Sigma Aldrich, 0.4 mM final), and ^13^C-, ^15^N-labeled lysine and arginine (Cambridge Isotope Laboratories, 0.4 mM final), respectively.

### Proteomics sample preparation for quantitative AP-MS (S-TRAP)

Quantitative AP-MS of AML1/ETO splice product with SILAC proteomics was performed with “heavy” and “light” labeled HEK293T cells stably expressing ‘AML1^N^-CAGE^N^’ and ‘CAGE^C^-ETO^C^’ constructs^55^. SILAC-labeled cells, grown to 70% confluency in two 10 cm cell plates each, were incubated with DMSO alone (light cells) or rapalog (100 nM, heavy cells) for 18 hours in SILAC DMEM. All cells were then washed with cold PBS (2 x) and lysed by following the subcellular fractionation protocol to isolate the nuclei. Then, each heavy and light isolated nuclei samples were benzonase nuclease digested by following the co-immunoprecipitation protocol described above. After combining fractions S1 and S2 of each sample, protein concentration was measured using the BCA assay (Thermo Fisher) and diluted with modified buffer A to 0.8 mg/mL protein concentration. For SILAC label-swap experiments, “light” HEK293T cells were incubated with rapalog, and “heavy” cells were incubated with DMSO and processed as above. 40 μL of anti-HA magnetic beads were pre-washed twice with modified buffer B and added per 1 mL sample (total 2 mL per condition) in a Lo-bind Eppendorf tube. The HA-IP was performed overnight with end-over-end rotation at 4 <C. Then, the beads were washed sequentially with 4x modified buffer B and 1x LC-MS grade H_2_O for 2 minutes each rotation. 100 μL of S-TRAP elution buffer (TEAB 50 mM, 5% SDS, pH 7.1) was added, and the beads were heated at 45 <C for 10 minutes. After collecting the eluent, beads were rinsed again with 100 μL of S-TRAP elution buffer and the eluants combined. Eluents from heavy (+) and light (-) co-immunoprecipitated samples were mixed 1:1 v/v ratio and processed for the S-TRAP sample preparation approach.

The SILAC mixture of co-enriched proteins in S-TRAP elution buffer was digested by following the manufacturer’s protocol (ProtiFi). In brief, proteins were reduced by TCEP (20 mM final) with orbital shaking for 10 minutes at 95 <C. After cooling to room temperature, samples were then alkylated by adding chloroacetamide (40 mM final) with orbital shaking in the dark for 30 minutes at room temperature. Aqueous phosphoric acid was added to a final concentration of ∼1.2% v/v. 1 volume of sample solution was then mixed with 6 volumes of S-TRAP binding buffer (90% methanol + 10% 1 M TEAB, pH 7.1) followed by centrifugation at 10,000g for 2.5 minutes. The supernatants were loaded on to an S-TRAP micro spin column and passed through by centrifugation (1,800 rpm, 1 minute) multiple times. The columns were then washed by adding S-TRAP binding buffer and centrifugation three times. Proteins captured on the columns were then digested by adding 20 μL of digestion buffer (50 mM TEAB, pH 8.5) containing 1 μg of sequencing grade trypsin (Promega) 1 hour at 47 <C. Following trypsinization, proteolyzed peptides were eluted by adding buffers and centrifuging (3,000g for 1 minute) sequentially with 40 μL digestion buffer, 40 μL of 0.2% formic acid in LC-MS grade water, 40 μL of formic acid in 40% acetonitrile, and 40 μL of formic acid in 60% acetonitrile. All eluted peptides were combined and dried using a SpeedVac (Thermo Fisher). Dried peptides were dissolved in 0.1% formic acid (pH 3) for proteomic analysis.

### Phosphopeptides enrichment from BCR/ABL or DNAJB1/PRKACA activated mammalian cells

Transfected HEK293T cells or stable AML12 cells were treated with rapalog (100 nM) for each desired timepoint; 0 minutes, 30-60 minutes, 120 minutes, 360 minutes, and 720 minutes (DNABJ1/PRKACA only). After finishing all treatments simultaneously, cells were washed twice with cold PBS and lysed in 1 mL of 8 M urea, 25 mM NH_4_HCO_3_ pH 8.0 supplement with EDTA-free Halt protease and phosphatase inhibitor cocktails per plate with sonication (Amplitude: 30%, No pulse, Time: 10 seconds x 2 times) at 4 <C. Lysates were centrifuged at 17,000 g for 10 minutes to remove insoluble material, and protein concentration was measured by BCA assay. Lysates were then diluted to 4 mg/mL protein concentration with 8 M Urea buffer, followed by disulfide reduction with dithiothreitol (DTT) (10 mM, 40 minutes, 50 <C), alkylation (iodoacetamide, 30 mM, 30 min, room temperature, protected from light) and quenching (DTT, 5mM, 10 minutes, room temperature). The proteome solution was diluted 4-fold with ammonium bicarbonate solution (25 mM, pH 8.0), CaCl_2_ added (1 mM), and digested with sequencing grade trypsin (∼1:100 enzyme/protein ratio; Thermo Fisher) at 37 <C while rotating overnight. Peptide digestion reactions were stopped by acidification to pH 2-3 with 2% formic acid, and peptides were then desalted with Sep-Pak tC18 cartridges (50-100 mg, Waters) and then lyophilized.

The lyophilized peptides from each condition were re-dissolved in Binding/Equilibration buffer (150 μL per 3 mg peptides) and applied to TiO_2_ spin tips (3 mg peptides per tip) for phosphopeptide enrichment according to the manufacturer’s protocol (High-Select^TM^ TiO_2_ Phosphopeptide Enrichment Kit, A32993, Thermo Scientific). Eluted phosphopeptides were transferred to Lo-bind Eppendorf tubes, and samples were immediately dried in a SpeedVac. To enrich any remaining phosphopeptides not captured using the TiO_2_ column the flow-throughs were passed over PHOS-Select IMAC (P9740, Sigma Aldrich) according to the manufacturer’s protocol. After drying the eluents, phosphopeptides from IMAC were combined with the TiO_2_-enriched peptides per fraction.

In the BCR/ABL studies, the enriched phosphopeptides were further used for immuno-affinity-based enrichment of phosphotyrosine peptides using a protocol adapted from reference^56^ with minor modifications, as described below. In brief, dried phosphopeptides from each fraction were re-dissolved in 800 μL cold IAP buffer (50 mM Tris-HCl, 150 mM NaCl, 1% w/v *n-*Octyl-β-D-glucopyranoside (EMD Millipore), EDTA-free Halt protease inhibitor and phosphatase inhibitor cocktails, pH 7.4). The peptide mixtures were agitated in a shaker for 30 minutes to dissolve the peptides thoroughly. Meanwhile, 200 μL of 25% anti-phosphotyrosine antibody (PY99) agarose beads (sc-7020 AC, Santa Cruz) were pre-washed twice with cold IAP buffer and added to the sample. The pY-IP was performed overnight with end-over-end rotation at 4 <C. After centrifugation (1,500g for 1 minute at 4 <C), the beads were washed sequentially with 3x IAP buffer and 2x LC-MS grade H_2_O for 2 minutes each rotation. 200 μL of 0.15% trifluoroacetic acid (TFA) in H_2_O was added, and the beads were agitated on a shaker for 15 minutes at room temperature to elute the phosphotyrosine peptides. Elution was repeated three times, and eluents were combined and desalted with ZipTip C18 tips (100 µL, Millipore) and dried in a SpeedVac.

### TMT labeling of enriched phosphopeptides

Phosphopeptides from different timepoint fractions (three biological replicates) were separately labeled with 16-plex isobaric tandem mass tags (TMTpro^TM^ 16plex Label Reagent Set, A44521, Thermo Fisher) according to the manufacturer’s protocol. In brief, 0.5 mg of TMTpro label reagents were equilibrated to room temperature, reconstituted with 20 µL of anhydrous acetonitrile (Optima grade), and added to dried peptides dissolved in 100 µL of TEAB (100 mM, pH 8.5). Labeling was carried out at room temperature for 2 hours with end-to-end rotation and quenched with 5 µL of 5% hydroxylamine (Sigma Aldrich) for 15 minutes. Finally, labeled peptides were combined into a new protein LoBind tube (Eppendorf) and dried in a SpeedVac before processing by the Princeton Proteomics and Mass Spectrometry Core Facility.

After drying, TMT-labeled peptides were re-dissolved in 300 µL of 0.1% TFA in water and fractionated into 8 fractions using the High pH Reversed-Phase Peptide Fractionation Kit (84868, Pierce). Fractions 1, 4, and 7 were combined as sample 1. Fractions 2 and 6 were combined as sample 2. Fractions 3, 5, and 8 were combined as sample 3. Three combined samples were dried in a SpeedVac and resuspended in 5% acetonitrile/water (0.1% formic acid, pH 3) for proteomics analysis.

### Proteomic LC-MS/MS runs

Proteomics samples were injected on to an Easy-nLC 1200 UPLC system employed a 45 cm x 100 μm (inner diameter) nano-capillary column packed with 1.9 μm C18-AQ resin (Dr. Maisch, Germany) mated to a metal emitter in-line with an Orbitrap Fusion Lumos (Thermo Scientific, USA). The column temperature was 45 <C, and a two-hour gradient method (SILAC) and a three-hour gradient method (TMT) were employed at a flow rate of 360 nL/min.

For running SILAC samples, the mass spectrometer was operated in data-dependent mode with a 120,000 resolution MS1 scan (positive mode, profile mode, AGC target of 4 x 10^5^, maximum IT of 50 ms, scan range of 375-1500 m/z) in the Orbitrap, followed by HCD fragmentation in the ion trap with 35% collision energy. A dynamic exclusion list was invoked to exclude previously fragmented peptides for 60 s, and a maximum cycle time of 3 s was used. The isolation window for precursor ions was set to 1.6 m/z in the quadrupole. The ion trap was operated in Rapid mode with an AGC target of 1 x 10^4^ and a maximum IT of 54 ms.

For running TMT labeled samples, the mass spectrometer was operated in data-dependent mode with synchronous precursor selection (SPS) – MS3 method with a 120,000 resolution MS1 scan (positive mode, profile mode, maximum IT of 5 x 10^3^, scan range of 400-1500 m/z) in the Orbitrap followed by CID fragmentation in the ion trap with 30% collision energy for MS2 and HCD fragmentation in the Orbitrap (50,000 resolution) with 55% collision energy for MS3. The MS3 scan range was set to 110-500 with an injection time of 200 ms. A dynamic exclusion list was invoked to exclude previously sequenced peptides for 60 s, and a maximum cycle time of 2.5 s was used. The isolation window for precursor ions was set to 0.7 m/z in the quadrupole. The ion trap was operated in Turbo mode.

### Proteomics data processing and analysis – AML1/ETO (SILAC qAP-MS, MaxQuant)

Data processing was performed using MaxQuant (v.2.3.1.0) software with default parameters unless otherwise noted. Replicates were analyzed in parallel with the match between runs selected. The UniProt human proteome reference database was used (proteome ID UP000005640), and Trypsin/P was selected as the enzyme. Methionine oxidation (+15.9949) and N-terminal acetylation (+42.0106) were specified as variable modifications and cysteine carbamidomethylation (+57.02146) as a static modification. For protein quantification, re-quantify was enabled, and the minimum label ratio count was set to one. Only proteins quantified in both experimental replicates were reported. Contaminants, including mitochondrial proteins, were excluded. Final outputs were filtered to achieve a < 1% false discovery rate (FDR).

The SILAC ratios of each protein from two biologically independent experiments were combined and converted to Log_2_ values, and *P* values were calculated by a univariate two-sided t-test and converted to -Log_10_ values. The waterfall plot was plotted with the median Log_2_ SILAC ratios of each protein. The proteins that passed the filter (rapalog-to-DMSO ratio > 2, *P* < 0.005) were considered significantly enriched proteins with the affinity pulldown of AML1/ETO splice product. Data are summarized in Supplementary Table 6.

### Proteomic data processing – BCR/ABL (TMT tyrosine phosphoproteomics, MaxQuant)

Data processing was performed using MaxQuant (v.2.3.1.0) software with default parameters unless otherwise noted. ‘Reporter ion MS3’ was chosen as a type for quantification, data were searched against the UniProt human proteome reference database (proteome ID UP000005640), and Trypsin/P was selected as the enzyme with a maximum of 3 missed cleavages. For processing TMTpro (16-plex) labeled samples in MaxQuant, parameters were imported with the correction of reporter ion isotopic distribution values provided from the TMTpro reagent set (Lot# WK334325). Each TMT 12-plex experiment was loaded as three fractions and three technical repeat runs. Methionine oxidation (+15.9949) and phosphorylation (+79.9663) on serine, threonine, and tyrosine were specified as variable modifications (maximum modification in the peptide set to 3), and cysteine carbamidomethylation (+57.02146) as a static modification. The maximum peptide mass was set to 6,000 Da, and the minimum label ratio count was set to one and quantified using both unique and razor peptides. Final outputs were filtered to achieve a < 1% false discovery rate (FDR).

### Proteomic data processing – DNAJB1/PRKACA (TMT phosphoproteomics, Proteome Discoverer)

The data were processed using Proteome Discoverer (v.2.5.0.400) software (Thermo Fisher) with customized workflows modified from a default version processing workflow ‘PWF_Tribird_TMTpro_Quan_SPS_MS3_SequestHT_Percolator’ and a consensus workflow ‘CWF_Comprehensive_Enhanced Annotation_Reporter_Quan.’ For searching data, the UniProt human (proteome ID UP000005640) or mouse (proteome ID UP000000589) proteome reference database was used with default parameters of the workflows unless otherwise noted below, and reporter ion isotropic distribution values were obtained from the TMTpro reagent set (Lot# XE341501). In the processing workflow, trypsin was selected as the enzyme with a maximum of 2 missed cleavages, and variable modifications of methionine oxidation (+15.995) and phosphorylation (+79.966) on serine, threonine, and tyrosine were specified. Cysteine carbamidomethylation (+57.021) and TMTpro reagents labeling on the N-terminus and lysine (+304.207) were set as static modifications. Site localization probability values were calculated using the ‘IMP-ptmRS’ node. In the consensus workflow, the site probability threshold was set to 70 in the ‘PSM Grouper’ node. The reporter co-isolation threshold was set to 90 in the ‘Reporter Ions Quantifier’ node, and motif information was obtained by adding the ‘Modification Sites’ node. Final outputs were filtered to achieve a <1% false discovery rate (FDR).

### Quantitative phosphoproteomics data analysis

The reporter ion intensities of identified phosphotyrosine peptides (BCR/ABL) or phosphopeptides (DNAJB1/PRKACA) were added up for each timepoint fraction from all LC-MS/MS runs, followed by consolidation with the same phosphosites (Supplementary Fig. 2b). Relative phosphorylation levels were calculated using the maximum TMT channel intensity, set to 100, within each protein phosphosite. Only phosphorylation dynamics of the phosphosites were counted for subsequent analyses when satisfying; (i) non-zero TMT ion intensities shown in at least 2 timepoint fractions in each biological replicate and (ii) notable changes between the values at a specific timepoint having average maximum value and the values at 0 minutes (*P* < 0.1 for BCR/ABL and *P* < 0.06 for DNAJB1/PRKACA, calculated by a univariate two-sided t-test). Data are summarized in Supplementary Table 7 (BCR/ABL, HEK293T), Supplementary Table 8 (DNAJB1/PRKACA, HEK293T), and Supplementary Table 9 (DNAJB1/PRKACA, AML12).

For clustering the protein phosphosites based on their phosphorylation dynamics, relative phosphorylation levels calculated by the method above were re-normalized to fold changes by the average value at 0 hours, and these re-normalized values were then converted to log_2_ values. The maximum fold change allowed in the dataset is set to 20 before log_2_ conversion. Then, phosphorylation sites were grouped statically and visualized as a heatmap using the Python and Seaborn package (v.0.11.2). Dynamic phosphorylation curves of phosphosites were plotted using the Prism software by the fold changes. Gene ontology analysis was performed by uploading protein lists to the Metascape website (https://metascape.org) and performing enrichment analysis^57^. Enriched sequence motifs of phosphosites were analyzed by uploading peptide sequences to the pLogo generator (https://plogo.uconn.edu) and performing enrichment analysis^58^. pSer/Thr sites identified in DNAJB1/PRKACA datasets (Supplementary Table 10, 11) were further analyzed by the Kinase Library platform^43^ to predict the clustered phosphorylation-site motifs and the kinase family targets (https://kinase-library.phosphosite.org/site), and if more than 4 kinases scored higher than 99%, the top 4 ranked candidates from each substrate prediction result were considered. Principal component analysis (PCA) was performed using the R package (BiocGenerics) with the log_2_ average fold-change values in each time point of the proteins in HEK293T (*P* < 0.06, n = 801) and AML12 (*P* < 0.06, n = 602) as inputs.

### Immunoprecipitation of AML1/ETO splice product from mammalian cells

10 cm of HEK293T cells transfected with plasmids encoding ‘FLAG-AML^N^-CAGE^N^’ and ‘CAGE^C^-ETO^C^’ were treated with rapalog (100 nM) for 18 hours. Cells were washed twice with cold PBS and lysed in 350 μL of 8 M urea, 25 mM NH_4_HCO_3_ pH 8.0 supplement with Roche cOmplete EDTA-free protease inhibitors. After sonication (Amplitude: 35%, Pulse: 1 second on / 1 second off, Time: 15 seconds) at 4 <C, lysates were centrifuged at 17,000 g for 10 minutes to remove insoluble material and then diluted ten times with 25 mM NH_4_HCO_3_ to 0.8 M Urea. 150 μL of 100% Anti-DYKDDDDK G1 affinity resin (GenScript) was prewashed with cold PBS (3 x) and then added to the 11 mL of lysates for FLAG-IP with end-over-end rotation at 4 <C overnight. After removing supernatants, the resins were washed sequentially with 8x 0.8 M Urea in PBS and 2x PBS. 250 μL of 2X Laemmli buffer supplement with 5 mM of DTT was added, and the beads were heated at 95 <C for 10 minutes, and the eluent collected. The splice product AML1/ETO was separated by 8% Bis-Tris SDS-PAGE gel, which was then stained with Coomassie stain until the band (∼100 kDa) was just visible. The product bands were isolated in gel pieces and subjected to in-gel reduction, alkylation, and overnight LysC digestion, as previously reported^59^. Peptides eluted from in-gel tryptic digestions were dried in a SpeedVac and were dissolved in 5% acetonitrile/water (0.1% formic acid, pH 3) for proteomics analysis (below).

### Direct MS1 and MS2 analysis of splice junction peptide in AML1/ETO splice product

Tryptic peptide of the splice junction in AML1/ETO splice product was initially validated by manual inspection of chromatograms and MS1 spectra with theoretical m/z values of peptide candidates; *m/z* = 519.517 (*z* = 4) and *m/z* = 692.353 (*z* = 3)

Validated MS1 ions were further analyzed by “MS2 run only” mode to identify the splice junction peptide with the desired b- and y-ions. The LC-MS/MS experiment was performed with an Easy-nLC 1200 UPLC system employing a 45 cm x 75 μm (inner diameter) nano-capillary column packed with 1.9 μm C18-AQ resin (Dr. Maisch, Germany) mated to a metal emitter in-line with an Orbitrap Fusion Lumos (Thermo Scientific, USA). The column temperature was 45 <C, and a one-hour gradient method was run at a flow rate of 300 nL/min. The mass spectrometer was operated to scan MS2 from MS1 of the inclusion list with a 60,000 resolution (positive mode, profile mode, AGC target of 4 x 10^5^, maximum IT of 118 ms) in the Orbitrap, followed by HCD fragmentation in the ion trap with 30% collision energy. The isolation window for precursor ions was set to 0.4 m/z in the quadrupole.

Raw files were processed with Thermo XCalibur Qual Browser, and MS2 peaks were manually annotated.

### Clonogenic assay of cells

Non-transformed and DNAJB1/PRKACA-expressing AML12 cells were seeded at 1,000 cells/well in 12-well plates. DMSO or GS-9007 (5 μM) was added to normal growth media the following day for inhibitor tests. Media with the drug was refreshed every 4 days. After 10 days of treatment, equivalent to at least 6 potential cell divisions, cells were rinsed in PBS and fixed in 4% paraformaldehyde in PBS for 20 minutes. Next, cells were stained with 0.1% crystal violet in 10% methanol, washed three times with water, and dried at room temperature for image capture using HP OfficeJet 3830 scanner. Finally, colonies were quantified using ImageJ with masking (convert to a binary image first, then *Image > Adjust > Threshold*) and particle analysis (*Analyze > Analyze particles*) tools.

---

**Extended Data Fig. 1.**
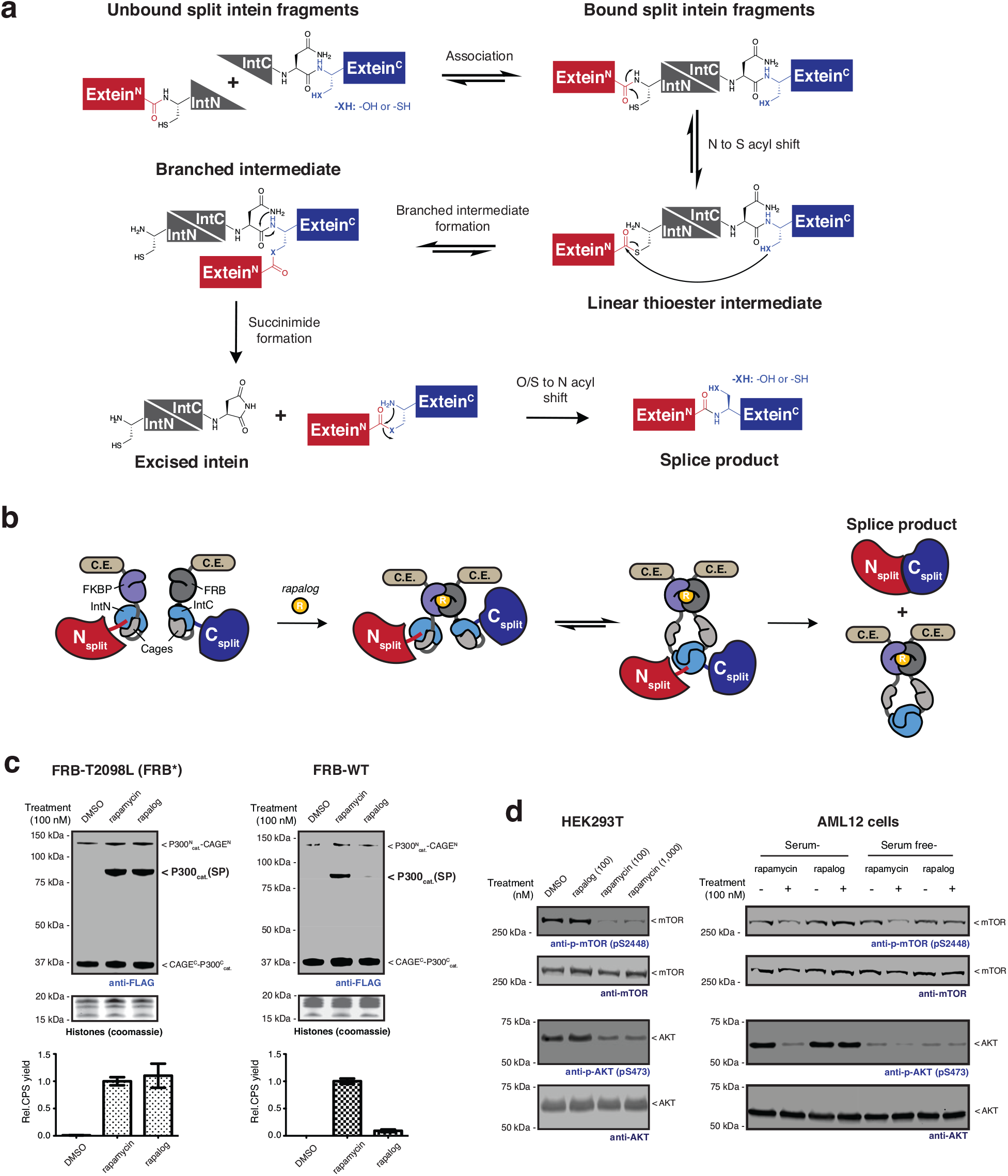
CAGE modules generate the splice products in live cells by non-toxic rapamycin analog treatment. **a,** Mechanism of the protein trans-splicing reaction by split inteins. **b,** Schematic of the CPS reaction using CAGE modules. Rapalog-induced FKBP-FRB dimerization is followed by a domain swapping event that yields an active split intein complex. **c,** Left – Immunoblots (top) of splice product (FLAG-tag) generation via CAGE modules incorporating FRB T2098L mutant (FRB*) in HEK293T cells. Cells were treated with DMSO, rapamycin (100 nM), or non-toxic rapamycin analog AP21967 (rapalog, 100 nM) for 18 hours. The bar graph below shows quantified splice product levels following the indicated treatments normalized to the product band intensity of rapamycin (bottom) (*n*=4, mean and s.e.m.). Right – as in the left-hand panel but employing wild-type FRB in the CAGE constructs. **d,** Immunoblots of phosphorylation levels at mTOR S2448 and AKT S473 sites in HEK293T cells (left) and AML12 cells (right) treated with DMSO, rapalog, or rapamycin at the indicated dose for 18 hours. Data in (**d**) is representative of *n*=3 independent experiments. CAGE^N^ refers to ‘NpuN^Cage^-FKBP,’ and CAGE^C^ refers to ‘FRB-NpuC^Cage^’ in (**c**).

**Extended Data Fig. 2.**
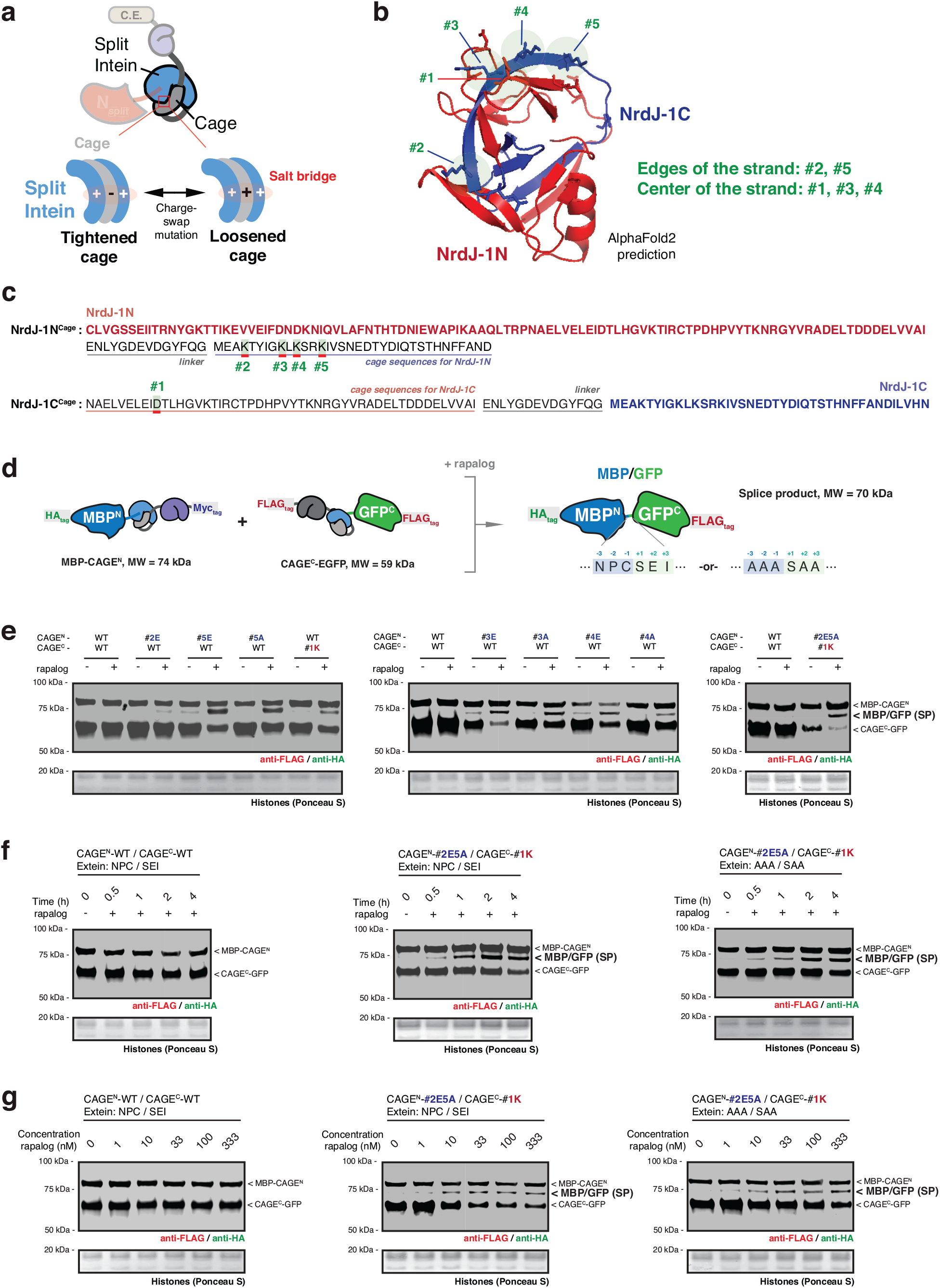
Engineering the CAGE toolbox via charge-swap mutations of the NrdJ-1 cages. **a,** Schematic showing engineering strategy used to tune the responsiveness of the CAGE modules via charge swap mutations. The interaction between the appended caging unit and the split intein can be strengthened or weakened by manipulating key salt bridges. **b,** A predicted structure of the NrdJ-1 intein generated via AlphaFold2 modeling, with the amino acids for charge-swap mutations in the cages marked in green circles. **c,** The amino acid sequences of both NrdJ-1N^Cage^ and NrdJ-1C^Cage^ constructs (intein fragment + appended cage) with the sites altered through charge-swap mutations indicated in (**b**) marked in green boxes, and underlined in red. **d,** Schematic of MBP-GFP splice product generated by proximity-triggered CPS between complementary MBP-CAGE^N^ and CAGE^C^-GFP constructs with rapalog treatment in cells. Molecular weights of the splice product and CAGE constructs are indicated. Flanking extein sequences at the splice site were either native NrdJ-1 (NPC/SEI) or AAA/SAA. **e,** Immunoblots of the MBP-GFP splice product (SP) bands in HEK293T cells co-expressing MBP-CAGE^N^ and CAGE^C^-GFP, treated with either DMSO or rapalog (100 nM) for 4 hours. Relative splice product levels were screened with every single mutant in the cages of the CAGE^N^ or CAGE^C^ indicated in (**b**-**c**) and the combination of #2E/#5A in the cage of CAGE^N^ and #1K in the cage of CAGE^C^. **f, g,** Immunoblots of the MBP-GFP splice product (SP) bands in HEK293T cells co-expressing MBP-CAGE^N^ and CAGE^C^-GFP, treated with either DMSO or rapalog (100 nM) for indicated time points (**f**) or treated with DMSO or increasing dose of rapalog for 4 hours (**g**). Left – CAGE without charge-swap mutations, NPC/SEI extein sequence. Middle – engineered CAGE (#2E/#5A in the cage of CAGE^N^ and #1K in the cage of CAGE^C^) NPC/SEI extein sequence. Right – engineered CAGE (#2E/#5A in the cage of CAGE^N^ and #1K in the cage of CAGE^C^), AAA/SAA extein sequence. Data in (**e**-**g**) are representative of *n*=3 independent experiments. CAGE^N^ refers to ‘NrdJ-1N^Cage^-FKBP’, and CAGE^C^ refers to ‘FRB-NrdJ-1C^Cage^’ in (**e**-**g**). Importantly, the engineered NrdJ-1 CAGE (#2E/#5A in the cage of CAGE^N^ and #1K in the cage of CAGE^C^) was found to be optimal and used in all subsequent studies.

**Extended Data Fig. 3.**
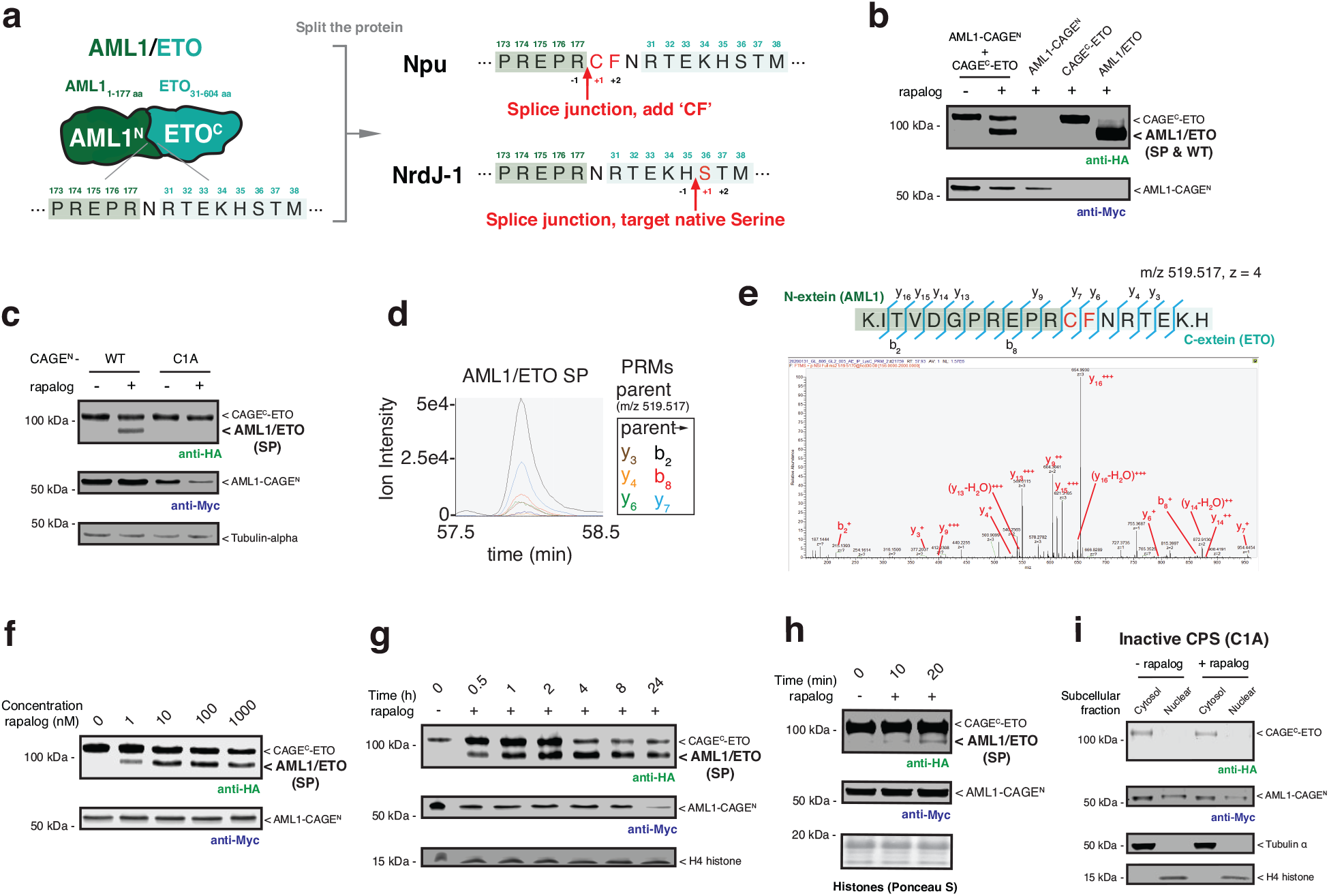
AML1/ETO splice product generated from co-expressed CAGE modules in cells. **a,** Protein splicing junctions employed in the generation of AML1/ETO using the caged Npu and NrdJ-1 split inteins. The caged Npu system required the insertion of a ‘CF’ di-peptide sequence to facilitate efficient protein trans-splicing. This ‘scar’ is added to the N-terminus of ETO segment. By contrast, the caged NrdJ-1 split intein supports traceless splicing using the native serine at position 36 of ETO. **b,** Immunoblots of HA-tagged AML1/ETO splice product (SP) generated in HEK293T cells co-expressing AML1-CAGE^N^ (Myc-tag) and CAGE^C^-ETO (HA-tag). Cells were treated with either DMSO or rapalog (100 nM) for 18 hours before analysis. HEK293T cells expressing a HA-tagged AML1/ETO fusion (WT) are used as a positive control (right most lane). **c,** Immunoblots of HA-tagged AML1/ETO splice product (SP) generated HEK293T cells co-expressing AML1-CAGE^N^ (wild-type or inactive C1A mutant) and CAGE^C^-ETO. Cells were treated with either DMSO or rapalog (100 nM) for 18 hours before analysis. **d, e,** Characterization of the AML1/ETO splice product by mass spectrometry. AML1/ETO splice product generated in HEK293T cells was subjected to trypsinolysis and targeted proteomics used to identify the tryptic peptide spanning the splice junction (Calculated *m/z* = 519.517, *z* = 4). Overlaid PRM chromatograms for the six parent-to-daughter transitions in targeted proteomics runs (**d**) and annotated MS/MS spectrum from the peptide (**e**). **f,** Immunoblots of AML1/ETO generation in HEK293T cells co-expressing AML1-CAGE^N^ and CAGE^C^-ETO. Cells were treated with DMSO or increasing doses of rapalog for 18 hours before analysis. **g, h,** Immunoblots of AML1/ETO generation in HEK293T cells co-expressing AML1-CAGE^N^ and CAGE^C^-ETO. Cells were treated with DMSO or rapalog (100 nM) for indicated time-points before analysis. **i,** Subcellular location of the AML1-CAGE^N^ (C1A mutant) and CAGE^C^-ETO in HEK293T cells treated with DMSO or rapalog (100 nM) for 18 hours before immunoblotting analysis. The C1A mutation of the CAGE^N^ module blocks the splice product generation. CAGE^N^ refers to ‘NrdJ-1N^Cage^-FKBP-NES’, and CAGE^C^ refers to ‘NES-FRB-NrdJ-1C^Cage^’. Data in (**b**-**c**) and (**f**-**i**) are representative of *n*=3 independent experiments. In panels (**b**-**h**), CAGE^N^ refers to ‘NpuN^Cage^-FKBP-NES,’ and CAGE^C^ refers to ‘NES-FRB-NpuC^Cage^.

**Extended Data Fig. 4.**
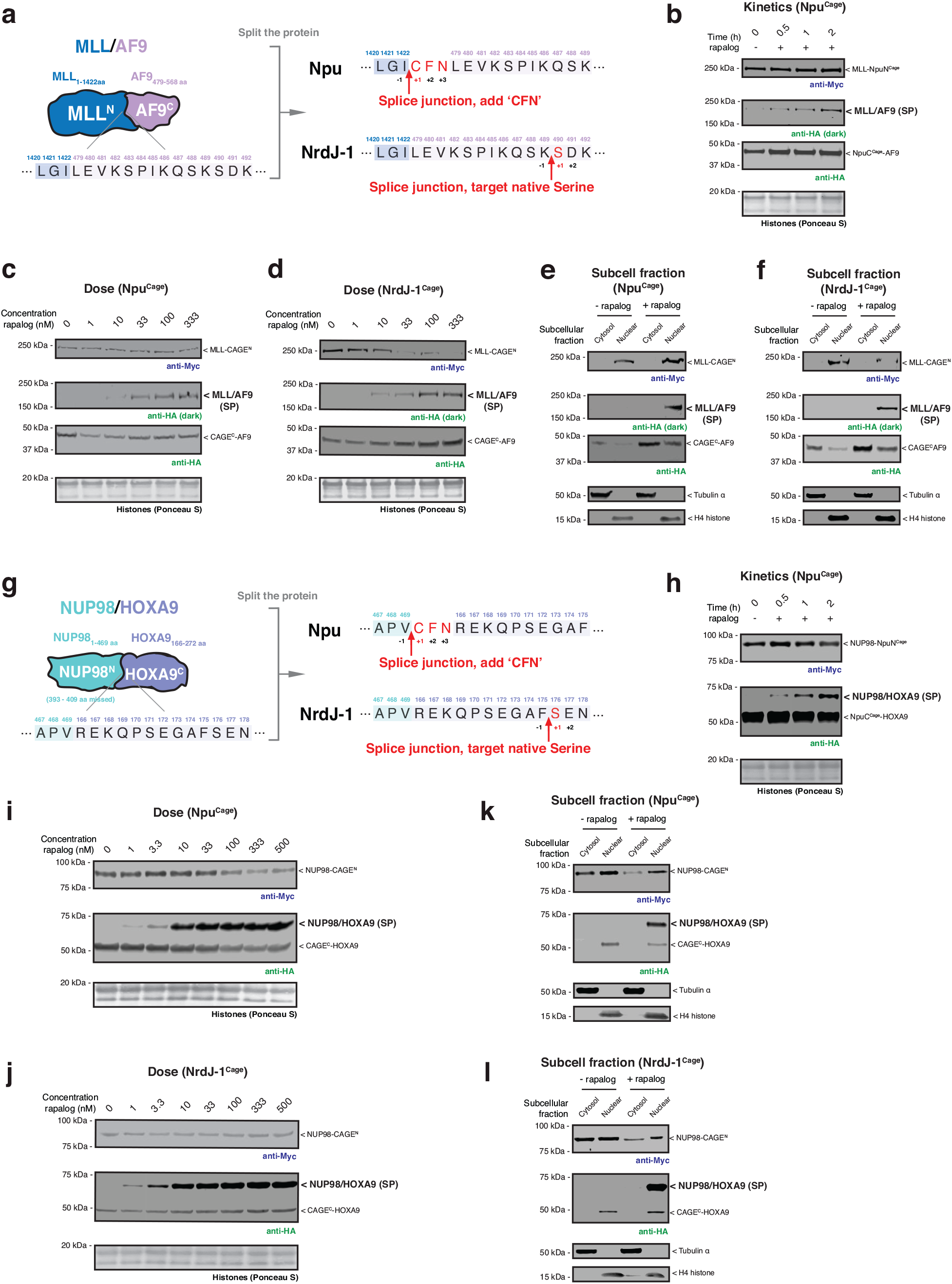
MLL/AF9 and NUP98/HOXA9 splice products generated from co-expressed CAGE modules in cells. **a,** Protein splicing junctions employed in the generation of MLL/AF9 using the caged Npu and NrdJ-1 split inteins. The caged Npu system required the insertion of a ‘CFN’ tri-peptide sequence to facilitate efficient protein trans-splicing. This ‘scar’ is added to the N-terminus of AF9 segment. The caged NrdJ-1 split intein supports traceless splicing using the native serine at position 490 of AF9. **b,** Immunoblots of MLL/AF9 (HA-tag) splice product (SP) generation in HEK293T cells co-expressing MLL-CAGE^N^ (Myc-tag) and CAGE^C^-AF9 (HA-tag). Cells were treated with DMSO or rapalog (100 nM) for indicated time-points before analysis. CAGE^N^ refers to ‘NpuN^Cage^-FKBP-NES,’ and CAGE^C^ refers to ‘NES-FRB-NpuC^Cage^.’ **c, d,** Immunoblots of MLL/AF9 generation in HEK293T cells co-expressing MLL-CAGE^N^ and CAGE^C^-AF9. Cells were treated with DMSO or increasing doses of rapalog for 18 hours before analysis. CAGE constructs incorporated either Npu^Cage^ (**c**) or NrdJ-1^Cage^ (**d**). **e, f,** Immunoblots of the HA-tagged MLL/AF9 splice product generated in HEK293T cells co-expressing MLL-CAGE^N^ and CAGE^C^-AF9 treated with rapalog (100 nM) for 18 hours followed by a subcellular fraction to separate cytosolic and nuclear proteins. Tubulin and histone H4 serve as cytosolic and nuclear markers, respectively. CAGE constructs incorporated either Npu^Cage^ (**e**) or NrdJ-1^Cage^ (**f**). **g,** Protein splicing junctions employed in the generation of NUP98/HOXA9 using the caged Npu and NrdJ-1 split inteins. The caged Npu system required the insertion of a ‘CFN’ tri-peptide sequence to facilitate efficient protein trans-splicing. This ‘scar’ is added to the N-terminus of HOXA9 segment. The caged NrdJ-1 split intein supports traceless splicing using the native serine at position 176 of HOXA9. **h,** Immunoblots of NUP98/HOXA9 (HA-tag) splice product (SP) generation in HEK293T cells co-expressing NUP98-CAGE^N^ (Myc-tag) and CAGE^C^-HOXA9 (HA-tag). Cells were treated with DMSO or rapalog (100 nM) for indicated time-points before analysis. CAGE^N^ refers to ‘NpuN^Cage^-FKBP-NES,’ and CAGE^C^ refers to ‘NES-FRB-NpuC^Cage^.’ **i, j,** Immunoblots of NUP98/HOXA9 generation in HEK293T cells co-expressing NUP98-CAGE^N^ and CAGE^C^-HOXA9. Cells were treated with DMSO or increasing doses of rapalog for 18 hours before analysis. CAGE constructs incorporated either Npu^Cage^ (**i**) or NrdJ-1^Cage^ (**j**). **k, l,** Immunoblots of the HA-tagged NUP98/HOXA9 splice product generated in HEK293T cells co-expressing NUP98-CAGE^N^ and CAGE^C^-HOXA9 treated with rapalog (100 nM) for 18 hours followed by a subcellular fraction to separate cytosolic and nuclear proteins. Tubulin and histone H4 serve as cytosolic and nuclear markers, respectively. CAGE constructs incorporated either Npu^Cage^ (**k**) or NrdJ-1^Cage^ (**l**). Data in (**b**-**f**) and (**h**-**l**) are representative of *n*=3 independent experiments.

**Extended Data Fig. 5.**
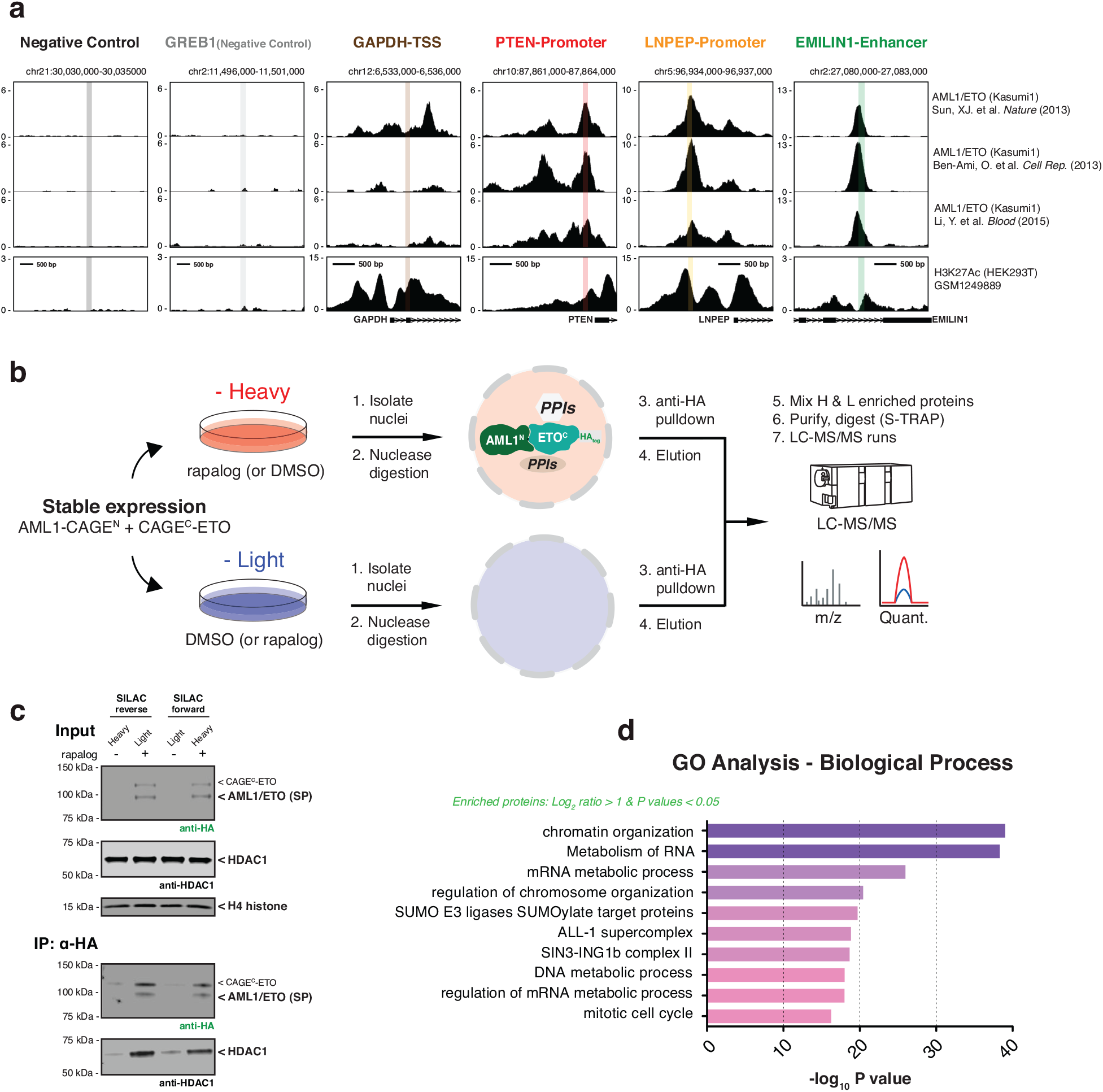
Multi-omics validation of the scarless AML1/ETO splice product. **a,** Screenshots of ChIP-seq results from three cited references displayed from the UCSC genome browser. Shown are six different regions with associated transcripts. The x-axis represents the genomic positions in base pairs: Negative Control_chr21:30,032,740-30,032,921; GREB1 (Negative Control)_chr2:11,498,624-11,498,837; GAPDH-TSS_chr12:6,534,775-6,534,872; PTEN-Promoter_chr10:87,863,242-87,863,355; LNPEP-Promoter_chr5:96,935,104-96,935,201; EMILIN1-Enhancer_chr2:27,081,454-27,081,577. The first two regions scored negative for AML1/ETO occupancy, and the last four scored positive for AML1/ETO occupancy in patient-derived Kasumi-1 cells. **b,** Schematic of the quantitative AP-MS workflow using SILAC-labeled HEK293T cells stably expressing AML1-CAGE^N^ and CAGE^C^-ETO constructs. **c,** Immunoblots of the abundance of AML1/ETO splice product (HA-tag) and HDAC1 in the SILAC inputs (top) and outputs (bottom). **d,** Gene Ontology (GO) biological process categories enriched among the proteins in quantitative AP-MS workflow, satisfying average SILAC ratio over two-fold and *P-*values less than 0.05. Statistical significance was determined by two-way student’s t-tests. CAGE^N^ refers to ‘NrdJ-1N^Cage^-FKBP-NES,’ and CAGE^C^ refers to ‘NES-FRB-NrdJ-1C^Cage^’ in (**c**).

**Extended Data Fig. 6.**
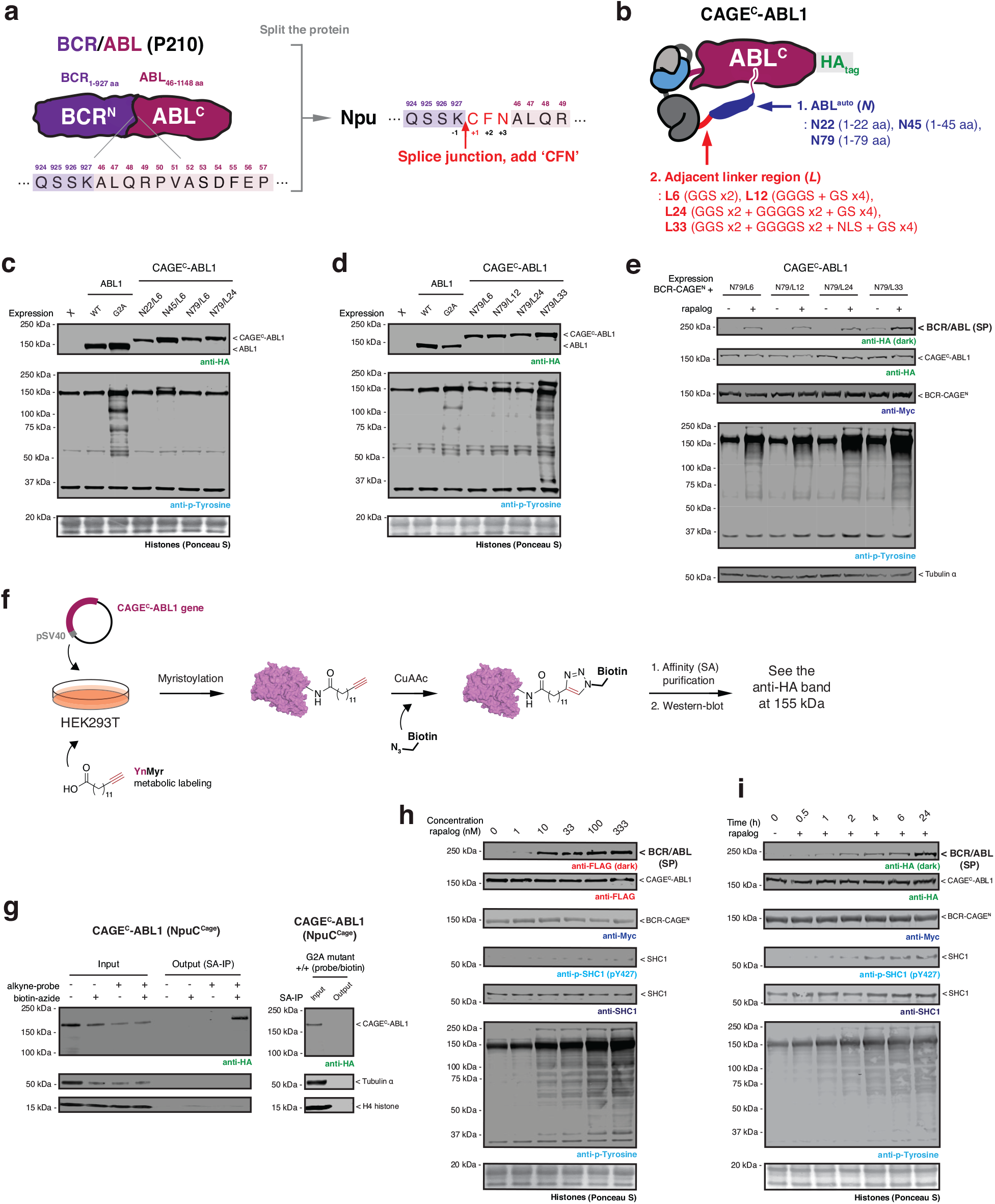
Design of CAGE^C^-ABL1 with appended native autoregulatory domain for inducible tyrosine phosphorylation in cells upon BCR/ABL activation. **a,** Schematic showing the design of the splicing site used for assembly of BCR/ABL using the CAGE system. ‘CFN’ amino acids are added to the N-terminus of the ABL segment to facilitate protein trans-splicing reaction using Npu^Cage^. **b,** Schematic showing the optimization of the autoinhibited CAGE^C^-ABL1 construct. Various lengths of the unstructured cap region containing the *N-*myristate site (Auto-regulatory domain, *N*) and the flexible linker connecting the auto-regulatory domain with the CAGE^C^ module (Adjacent linker region, *L*) were screened to find the optimal arrangement needed to suppress ABL kinase activity. **c, d,** Immunoblots of the basal tyrosine phosphorylation levels in HEK293T cells expressing indicated ABL constructs and CAGE^C^-ABL1 variants featuring different autoinhibitory domain (*N*) and adjacent linker (*L*) lengths. **e,** Immunoblots of BCR/ABL (HA-tag) and global tyrosine phosphorylation levels in HEK293T cells co-expressing BCR-CAGE^N^ and CAGE^C^-ABL1 variants. Cells treated with DMSO or rapalog (100 nM) for 18 hours before analysis. From these studies, it was determined that the optimal CAGE^C^-ABL1 construct contained a L79 + N24 combination. **f,** Schematic of the *N*-myristoylation metabolic labeling workflow. HEK293T cells expressing optimal CAGE^C^-ABL1 (N79/L24) were cultured in growth media with YnMyr (20 µM) for 24 hours and lysed. Labeled proteins were captured by click chemistry reaction with an azide-biotin tag, enriched with streptavidin beads, and analyzed by anti-HA immunoblots. **g,** Immunoblots of the HA-tagged CAGE^C^-ABL1 construct expressed in HEK293T cells cultured with YnMyr and enriched by the workflow described in (**f**). CAGE^C^-ABL1 construct was enriched following YnMyr metabolic labeling and biotin conjugation (left). By contrast, the G2A mutation of CAGE^C^-ABL1 failed to be enriched in the workflow (right). **h,** Proximity-induced CPS in HEK293T cells co-expressing BCR-CAGE^N^ (Myc-tag) and CAGE^C^-ABL1 (HA-tag). Cell were treated with DMSO or increasing doses of rapalog (100 nM) for 18 hours before immunoblotting using the indicated antibodies. **i,** Kinetics of BCR/ABL splice product generation in HEK293T cells co-expressing BCR-CAGE^N^ and CAGE^C^-ABL1. Cells were treated with DMSO or rapalog (100 nM) for indicated time-points, followed by immunoblotting using indicated antibodies. Data in (**c**-**e**) and (**g**-**i**) are representative of *n*=3 independent experiments. CAGE^N^ refers to ‘NpuN^Cage^-FKBP,’ and CAGE^C^ refers to ‘ABL^auto^-FRB-NpuC^Cage^’ in (**c**-**e**) and (**g**-**i**).

**Extended Data Fig. 7.**
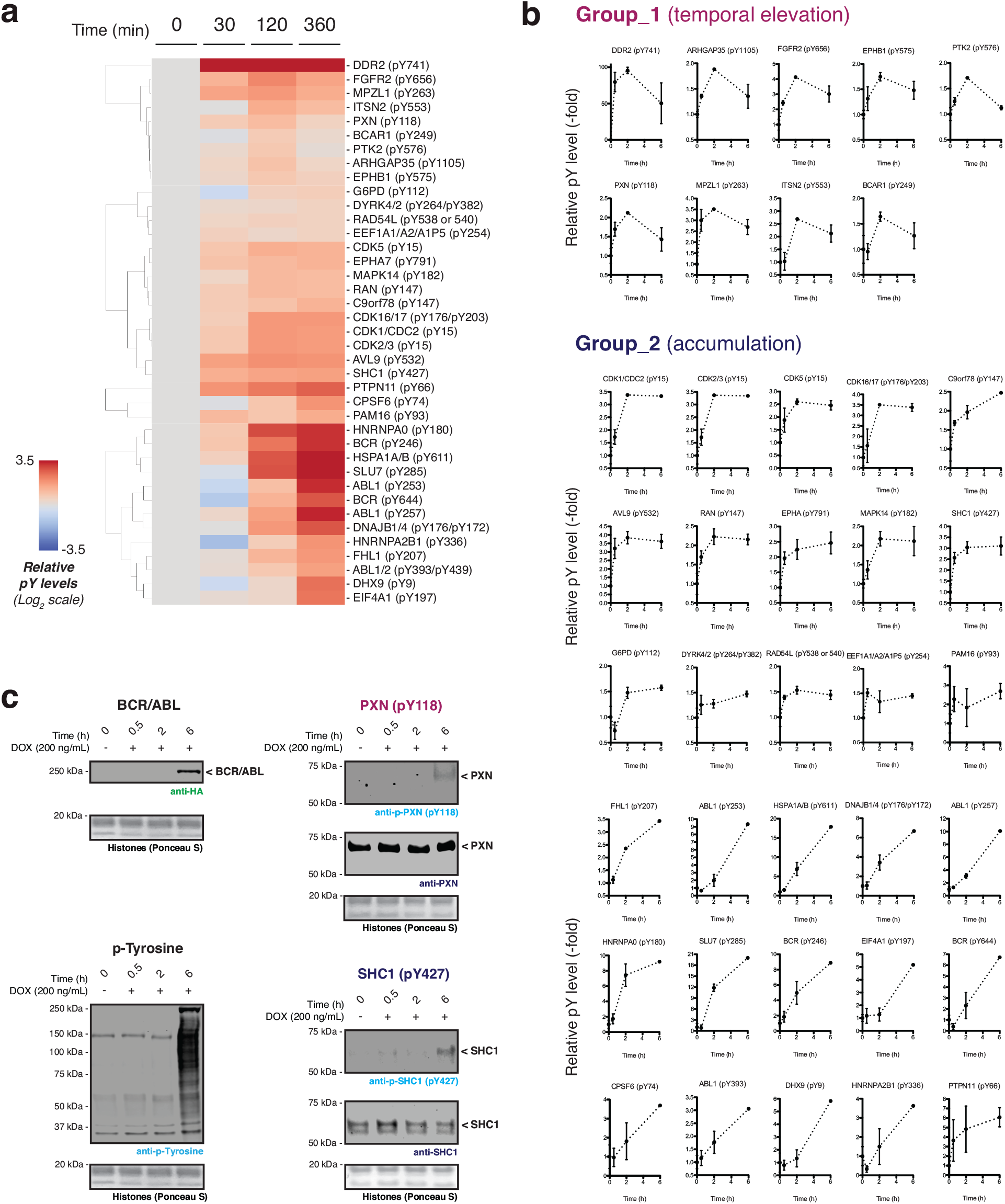
Dynamic tyrosine phosphorylation events following BCR/ABL activation. **a,** Cluster map of the proteomics dataset representing the time-resolved phosphorylation at protein tyrosine sites, combined from three biological replicates. Data processing employing the stringency filters described in fig. S8B results in 39 representative dynamics within 6-hour timeslots. **b,** Phosphorylation fold-change plots of representative protein tyrosine sites as a function of time. Phosphorylation levels at each time point are normalized to the values at 0 hours (*n=*3. Mean with s.e.m). **c,** Immunoblots of tyrosine phosphorylation levels in HEK293T cells containing a Dox-inducible BCR/ABL gene expression system. BCR/ABL expressed via doxycycline (200 ng/mL) treatment for indicated time-points. Data in (**a**-**b**) were generated from *n*=3 independent biological replicates. Data in (**c**) are representative of *n*=3 independent experiments. CAGE^N^ refers to ‘NpuN^Cage^-FKBP,’ and CAGE^C^ refers to ‘ABL^auto^-FRB-NpuC^Cage^’.

**Extended Data Fig. 8.**
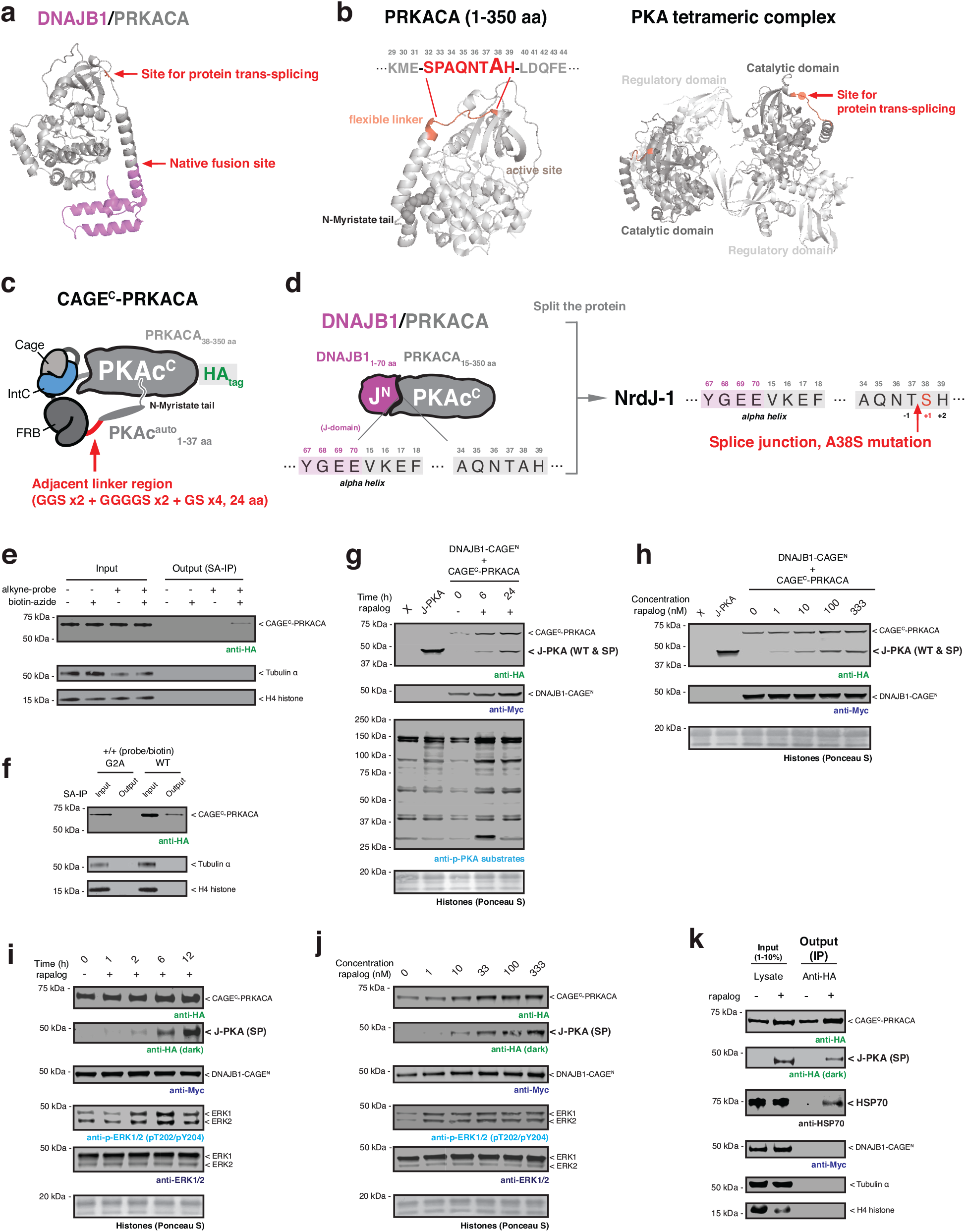
Design of DNAJB1-CAGE^N^ and CAGE^C^-PRKACA for inducible kinase activation in cells. **a,** Crystal structure of the DNAJB1/PRKACA fusion oncoprotein (PDB: 4WB7). The native gene fusion site and the potential ligation site for protein splicing are indicated. **b,** Left – crystal structure of mouse PRKACA (PDB: 4DFX) highlighting the myristate tail bound to the allosteric site in the catalytic domain. A flexible linker colored in red is a potential site for insertion of the CAGE^C^ module. Right – crystal structure of PKA tetrameric complex (PDB: 3TNP). The location of the CAGE^C^ module insertion site is indicated in red. **c,** Schematic depicting the design of the CAGE^C^-PRKACA construct. The N-terminal alpha-helix of PRKACA containing the *N-*myristoylation site (1-37 aa) is connected to the N-terminus of the CAGE^C^ module via a 24 amino acids flexible linker. **d,** Schematic of the split site used within DNAJB1/PRKACA incorporation of the CAGE modules. A38 amino acid site in the PRKACA is mutated to serine in the N-terminus of split PRKACA for protein trans-splicing reaction by NrdJ-1^Cage^. **e, f,** Immunoblots of the HA-tagged CAGE^C^-PRKACA construct expressed in HEK293T cells cultured with YnMyr and enriched using the workflow described in fig. S7F. CAGE^C^-PRKACA construct was enriched following YnMyr metabolic labeling and biotin conjugation (**e**). By contrast, the G2A mutation of CAGE^C^-PRKACA failed to be enriched in the workflow (**f**). **g,** Proximity-induced CPS in HEK293T cells co-expressing DNJAB1-CAGE^N^ (Myc-tag) and CAGE^C^-PRKACA (HA-tag). Cells were treated with DMSO or rapalog (100 nM) for indicated time-points before immunoblotting. X refers to untransfected cells, whereas J-PKA refers to cells expressing the positive control DNAJB1/PRKACA fusion (HA tagged). **h,** Similar to panel (**g**) but showing the dose-response to rapalog at the 24-hour timepoint. **i,** Proximity-induced CPS in AML12 cells co-expressing DNJAB1-CAGE^N^ (Myc-tag) and CAGE^C^-PRKACA (HA-tag). Cells were treated with DMSO or rapalog (100 nM) for indicated time-points before immunoblotting. **j,** Similar to panel (**i**) but showing the dose-response to rapalog at the 6-hour timepoint. **k,** Immunoblots of HSP70 co-immunoprecipitated with HA-tagged splice product. A pulldown assay was performed using the lysates from stable AML12 cells treated with DMSO or rapalog (100 nM) for 24 hours. Data in (**e**-**k**) are representative of *n*=3 independent experiments. CAGE^N^ refers to ‘NrdJ-1N^Cage^-FKBP,’ and CAGE^C^ refers to ‘PKAc^auto^-FRB-NrdJ-1C^Cage^’ in (**e**-**k**).

**Extended Data Fig. 9.**
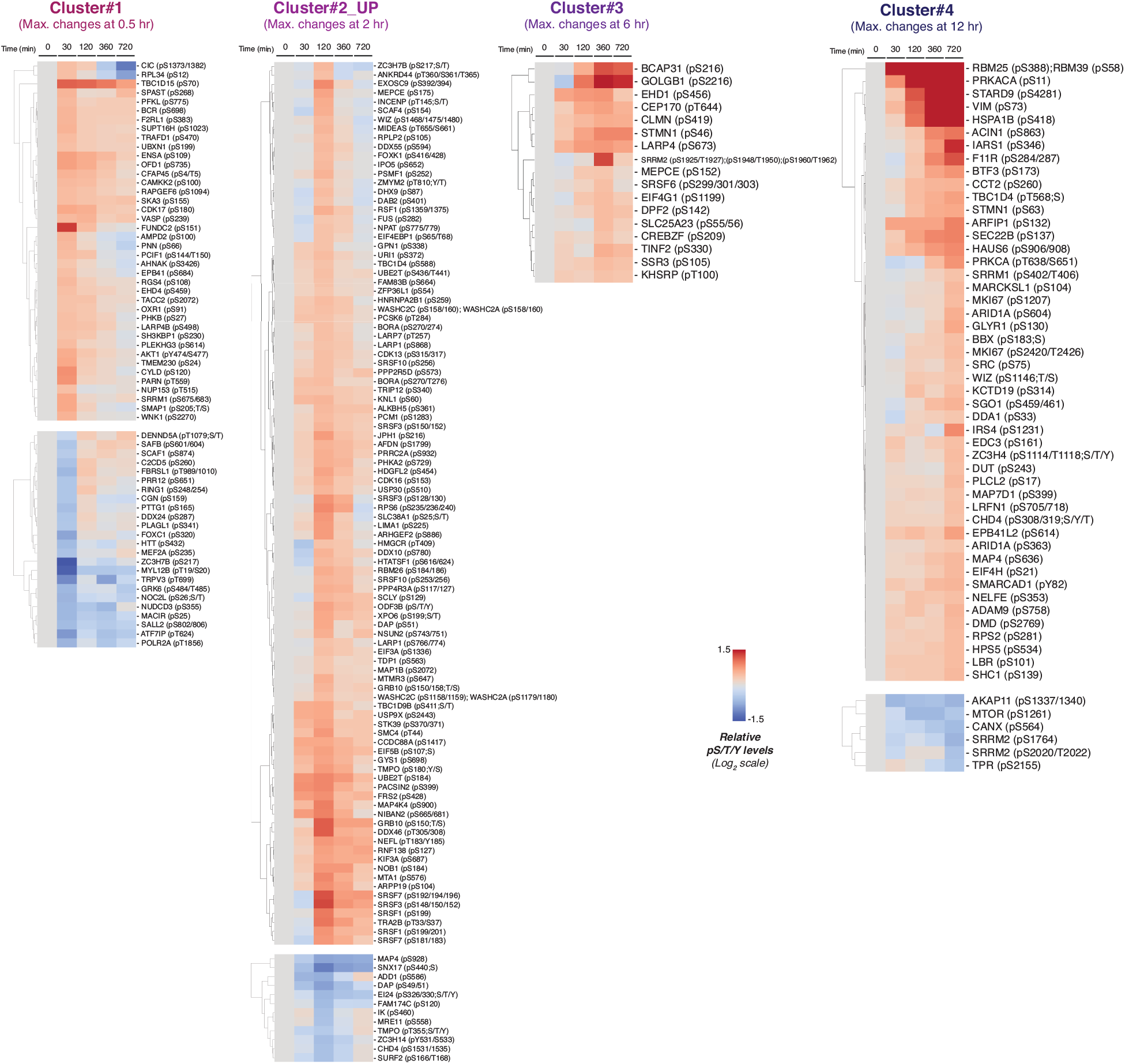
Clustering of the phosphorylation dynamics following DNAJB1/PRKACA generation in HEK293T cells. Protein phosphorylation dynamics in the HEK293T proteomics dataset are grouped into four clusters by the time of average maximum changes following induction of protein splicing. Relative phosphorylation levels in Cluster#1 members are maximally changed by 30 minutes, Cluster#2 members in 2 hours, Cluster#3 members in 6 hours, and Cluster#4 members in 12 hours. The members in each cluster were subcategorized into ‘UP- and DOWN-phosphorylated’ sites and proteins.

**Extended Data Fig. 10.**
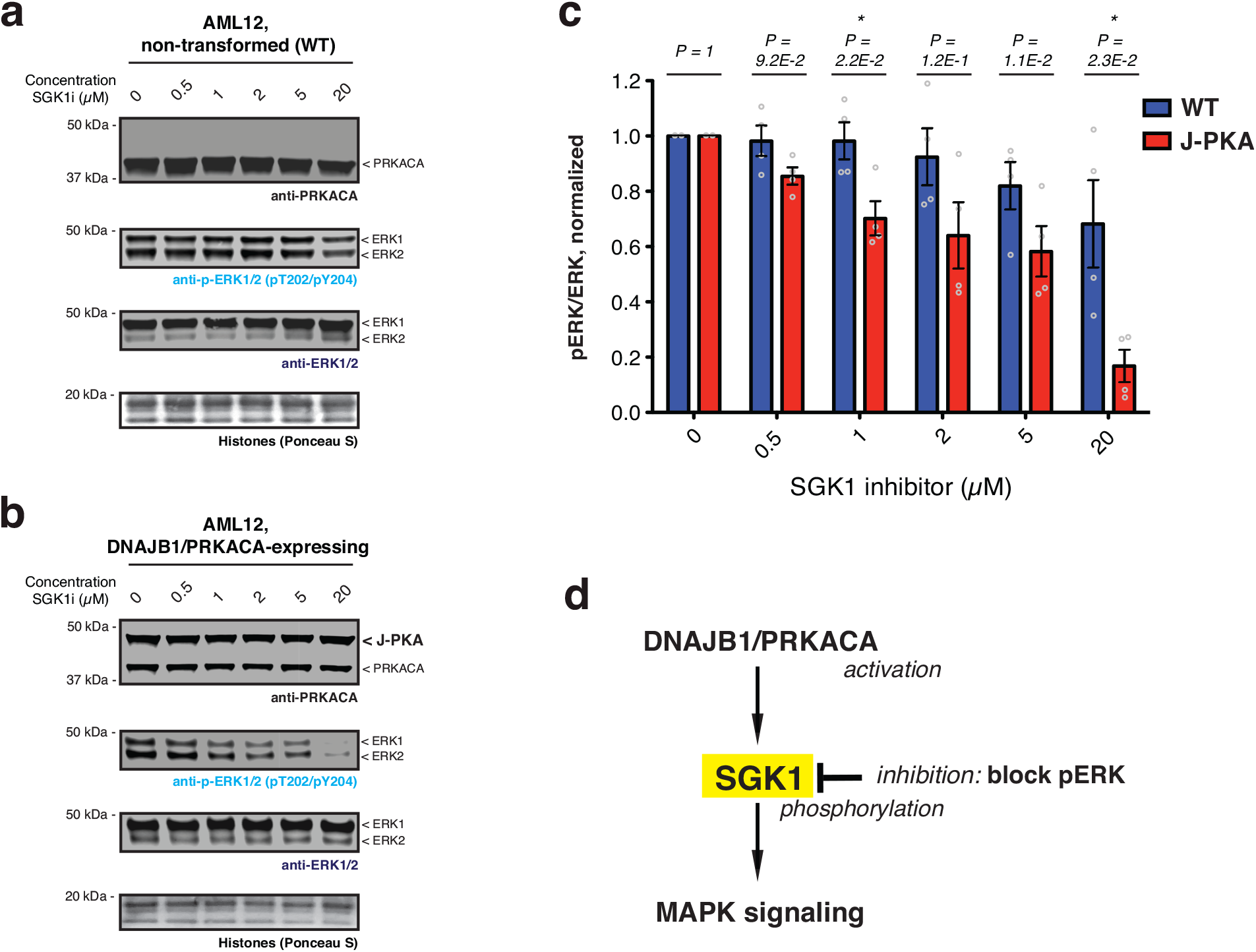
Pharmacologically inhibiting SGK1 in DNAJB1/PRKACA-expressing AML12 cells. **a, b,** Immunoblots of phospho-ERK1/2 in non-transformed (**a**) and DNAJB1/PRKACA-expressing (**b**) AML12 cells. Cells were treated with DMSO or increasing doses of GS-9007 for 24 hours followed by immunoblotting using the indicated antibodies. **c,** Bar charts showing quantified pERK levels of indicated treatments in non-transformed and DNAJB1/PRKACA-expressing AML12 cells normalized by the ERK intensity (*n*=4, means and s.e.m). Statistical significance was determined by two-way student’s t-tests: *P < 0.05; **P < 0.005; ***P < 0.001. **d,** Schematic of a proposed molecular link between DNAJB1/PRKACA activation and ERK phosphorylation through SGK1.

