## Supplementary Figures for "Distinct phases of cellular signaling revealed by time-resolved protein synthesis"

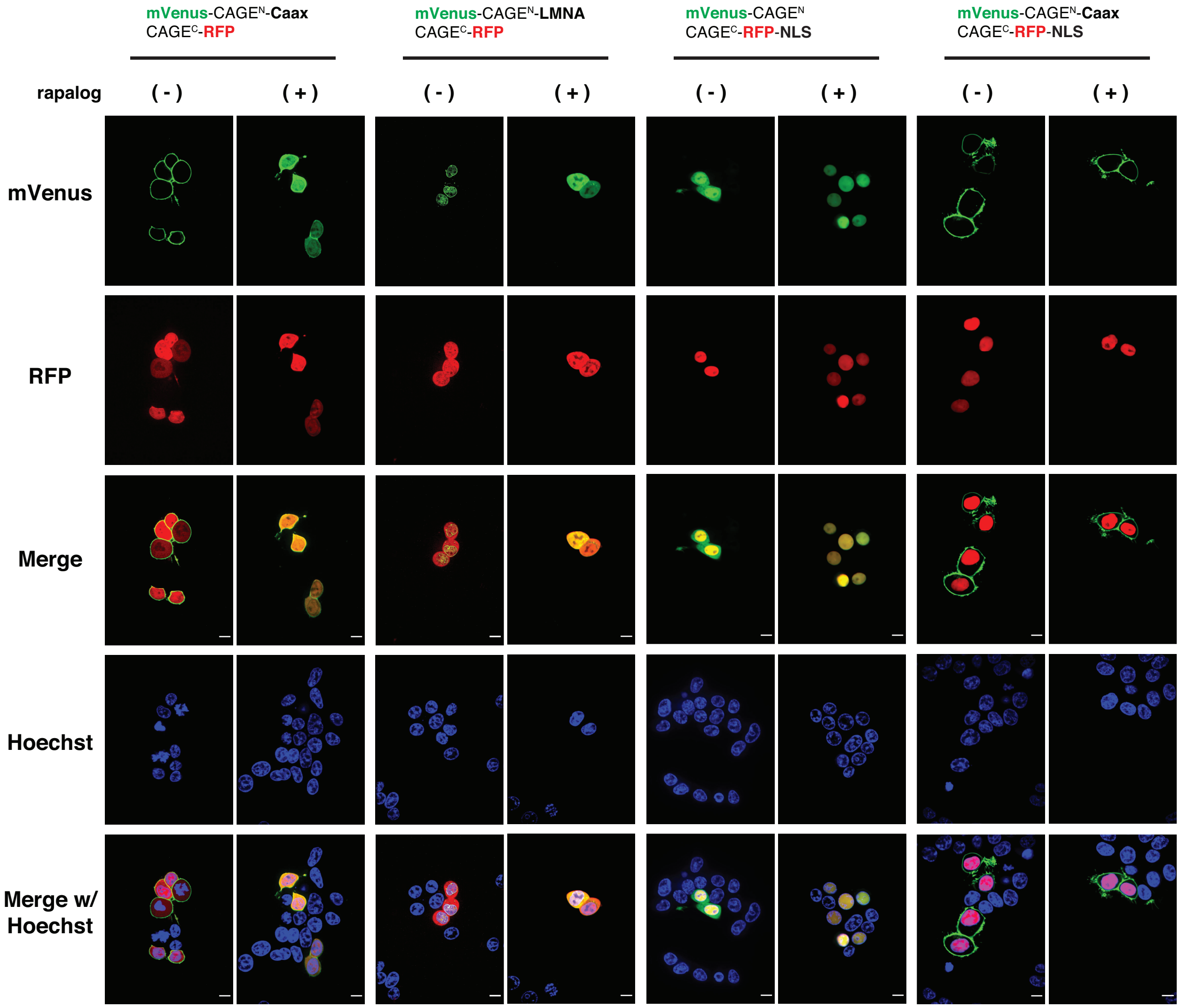


**Supplementary Fig. 1 | Uncropped fluorescence imaging of mVenus and RFP localization in live cells.** HEK293T cells co-expressing mVenus-CAGE$^{N}$ and CAGE$^{C}$-RFP were treated with either DMSO or rapalog (100 nM) for 10 hours. mVenus (green) and RFP (red) were visualized by their intrinsic fluorescence, and Hoechst 33342 (blue) was used as a nuclear marker. Scale bars, 10 μm. Full definition of control elements used: Caax sequence – KKKKKKSKTKCVIM (C-terminal fusion); LMNA – Prelamin-A/C (residues 2-664, C-terminal fusion); NLS sequence – PAAKRVKLD (c-Myc, C-terminal fusion).

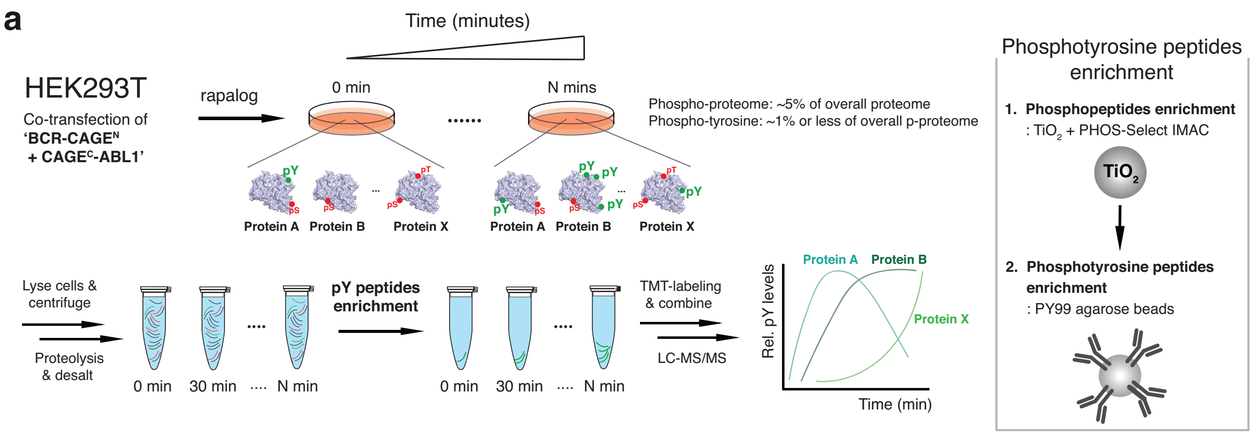


* **'DNAJB1-CAGE^N + CAGE^C-PRKACA'**
: Co-transfected HEK293T or stable AML12, Phosphopeptides enrichment by $TiO_2$ only (related to Fig. 4.)

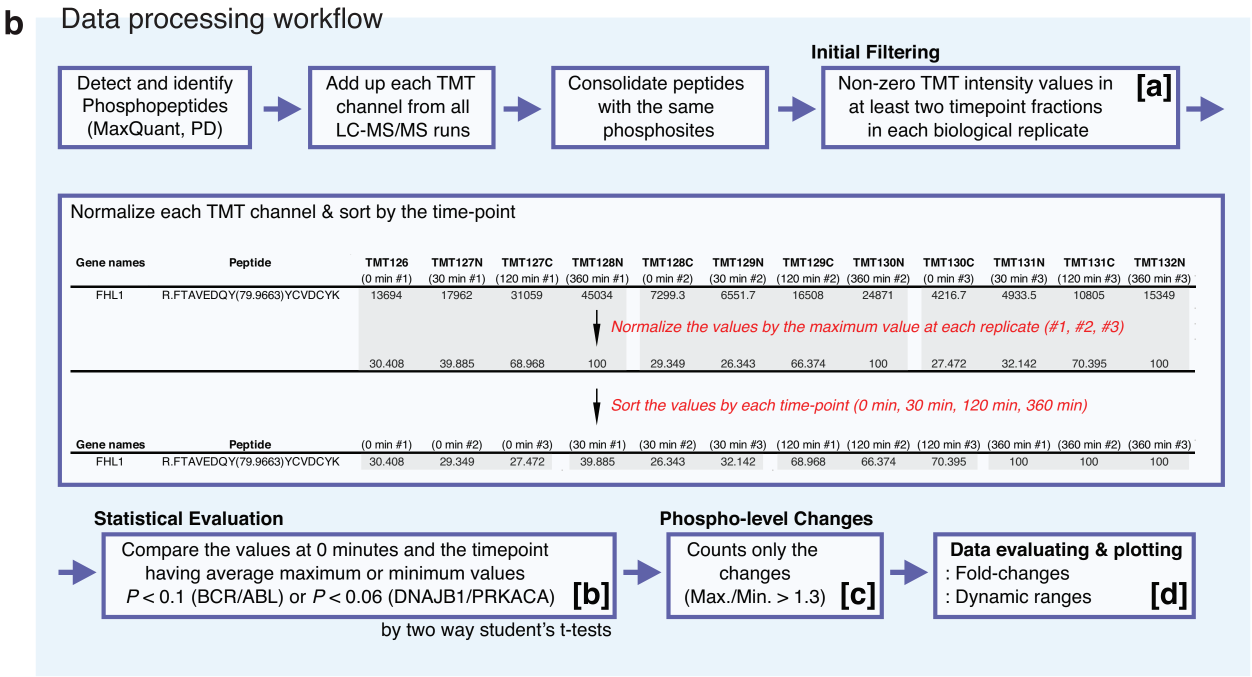


| Gene names | Peptide | TMT126 (0 min #1) | TMT127N (30 min #1) | TMT127C (120 min #1) | TMT128N (360 min #1) | TMT128C (0 min #2) | TMT129N (30 min #2) | TMT129C (120 min #2) | TMT130N (360 min #2) | TMT130C (0 min #3) | TMT131N (30 min #3) | TMT131C (120 min #3) | TMT132N (360 min #3) |
|---|---|---|---|---|---|---|---|---|---|---|---|---|---|
| FHL1 | R.FTAVEDQY(79.9663)YCVDCYK | 13694 | 17962 | 31059 | 45034 | 7299.3 | 6551.7 | 16508 | 24871 | 4216.7 | 4933.5 | 10805 | 15349 |
| | | 30.408 | 39.885 | 68.968 | 100 | 29.349 | 26.343 | 66.374 | 100 | 27.472 | 32.142 | 70.395 | 100 |

| Gene names | Peptide | (0 min #1) | (0 min #2) | (0 min #3) | (30 min #1) | (30 min #2) | (30 min #3) | (120 min #1) | (120 min #2) | (120 min #3) | (360 min #1) | (360 min #2) | (360 min #3) |
|---|---|---|---|---|---|---|---|---|---|---|---|---|---|
| FHL1 | R.FTAVEDQY(79.9663)YCVDCYK | 30.408 | 29.349 | 27.472 | 39.885 | 26.343 | 32.142 | 68.968 | 66.374 | 70.395 | 100 | 100 | 100 |

**c** # of phosphopeptides in the 'Data processing workflow'

| | 'BCR-CAGE^N + CAGE^C-ABL1' | 'DNAJB1-CAGE^N + CAGE^C-PRKACA' (related to Fig. 4.) | |
|---|---|---|---|
| | HEK293T | HEK293T | AML12 |
| [a] | 50 | 3740 | 3130 |
| [b] | 39 | 801 | 602 |
| [c] | 39 | 373 | 296 |
| [d] | 39 | 245 | 138 |

**Supplementary Fig. 2 | Time-resolved quantitative tyrosine phosphoproteomics workflow. a,** Schematic of the experimental workflow used to characterize changes in global pTyr phosphorylation levels following generation of BCR/ABL by CPS. HEK293T cells co-expressing BCR-CAGE$^N$ and CAGE$^C$-ABL1 constructs were treated with rapalog (100 nM) for desired time points. Cells were then lysed and the proteome trypsinized. Following a desalting step, phosphotyrosine peptides were enriched and labeled with unique tandem mass tag channels. The labeled samples were pooled and subjected to LC-MS/MS analysis. (Note, time-resolved phosphoproteomics downstream of DNAJB1/PRKACA activation [see Fig. 4.] was performed using a similar workflow, except the phosphopeptide enrichment step involved $TiO_2$ affinity capture only.) **b,** Schematic depicting the data processing workflow employed in phosphoproteomics. After searching for phosphopeptides (Peptide FDR < 0.01), the reporter ion intensities of detected peptides were combined and converted to the relative abundance to generate phosphorylation dynamic curves for each protein phosphorylation site. All data were generated from *n*=3 independent biological replicates. **c,** Number of phosphopeptides filtered in each step of the data processing workflow. The phosphopeptides having non-zero TMT ion intensity values in at least two timepoint fractions in each biological replicate are referred to as [a]. Within group [a], the phosphopeptides in [b] exhibit statistically significant changes between the values at a specific timepoint and those at 0 minutes, as determined by a univariate two-sided t-test. The significance thresholds used are $P < 0.1$ for BCR/ABL and $P < 0.06$ for DNAJB1/PRKACA. Furthermore, among the phosphopeptides in [b], those in [c] demonstrate notable phospho-level changes during the time course (Max./Min. > 1.3). Within set [c], group [d] represents the phosphopeptides selected for the phosphorylation dynamics analysis. For DNAJB1/PRKACA, its dynamics are determined based on the filtering criteria outlined in Supplementary Fig. 3a and Supplementary Fig. 4a showing either UP- or DOWN-phosphorylation.

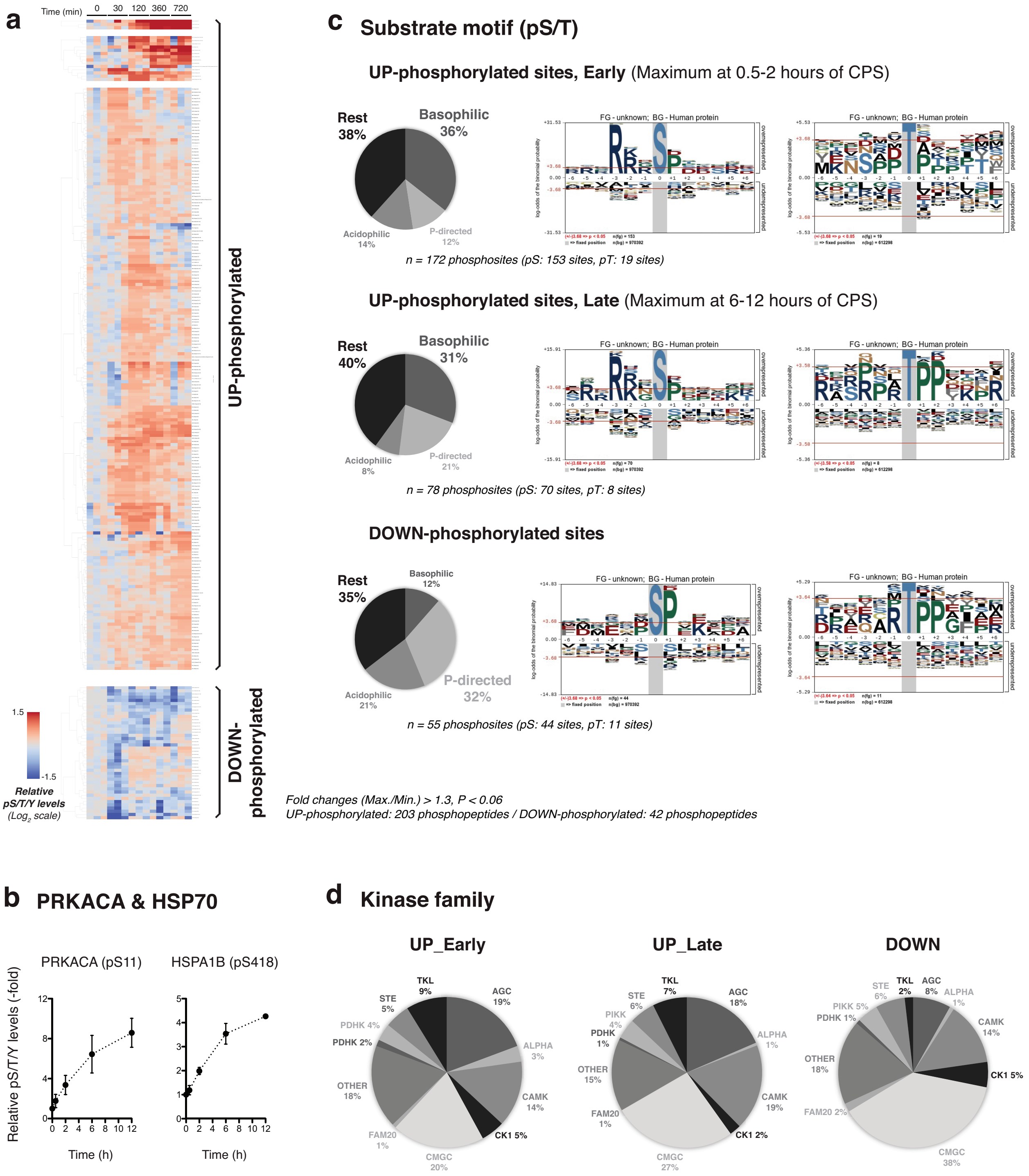


**Supplementary Fig. 3 | Dynamic protein phosphorylation in HEK293T cells following DNAJB1/PRKACA generation by CPS. a,** Cluster map of the proteomics dataset representing the time-resolved phosphorylation combined from three biological replicates. Data processing with the stringency filters described in Supplementary Fig. 2b results in 245 representative dynamics over the 12-hour time period. 'UP-phosphorylated' proteins corresponded to at least a 30% increase compared to the average phosphorylation levels at time zero (Max./0 hr value > 1.3-fold, n = 203). 'DOWN-phosphorylated' proteins corresponded to at least a 50% decrease from maximum to minimum and a 20% or more decrease from time zero to minimum average phosphorylation levels within the 12-hour timeslot (0 hr/Min. value > 1.2-fold & Max./Min. value > 1.5-fold, n = 42). **b,** Fold-change in phosphorylation of PRKACA and HSP70 at indicative times. Phosphorylation levels at each time point are normalized to the values at 0 hours (*n*=3. Mean with s.e.m). **c,** Pie charts (left) summarizing clustered phosphorylation-site motifs (predicted by The Kinase Library[43]) and enriched sequence motifs (right) derived from serine/threonine phosphorylation sites sorted by their dynamics. 'UP_Early' protein modification sites showed maximum phosphorylation at 0.5-2 hours of CPS, and 'UP_Late' member sites had maximum phosphorylation at 6-12 hours of CPS. 'DOWN' indicates the modification sites of the 'Down-phosphorylated' proteins in (**a**). **d,** Pie charts summarizing analysis of the three phosphorylation groups (UP_Early, UP_Late, DOWN) against the clustered human serine/threonine kinome (predicted by The Kinase Library[43]).

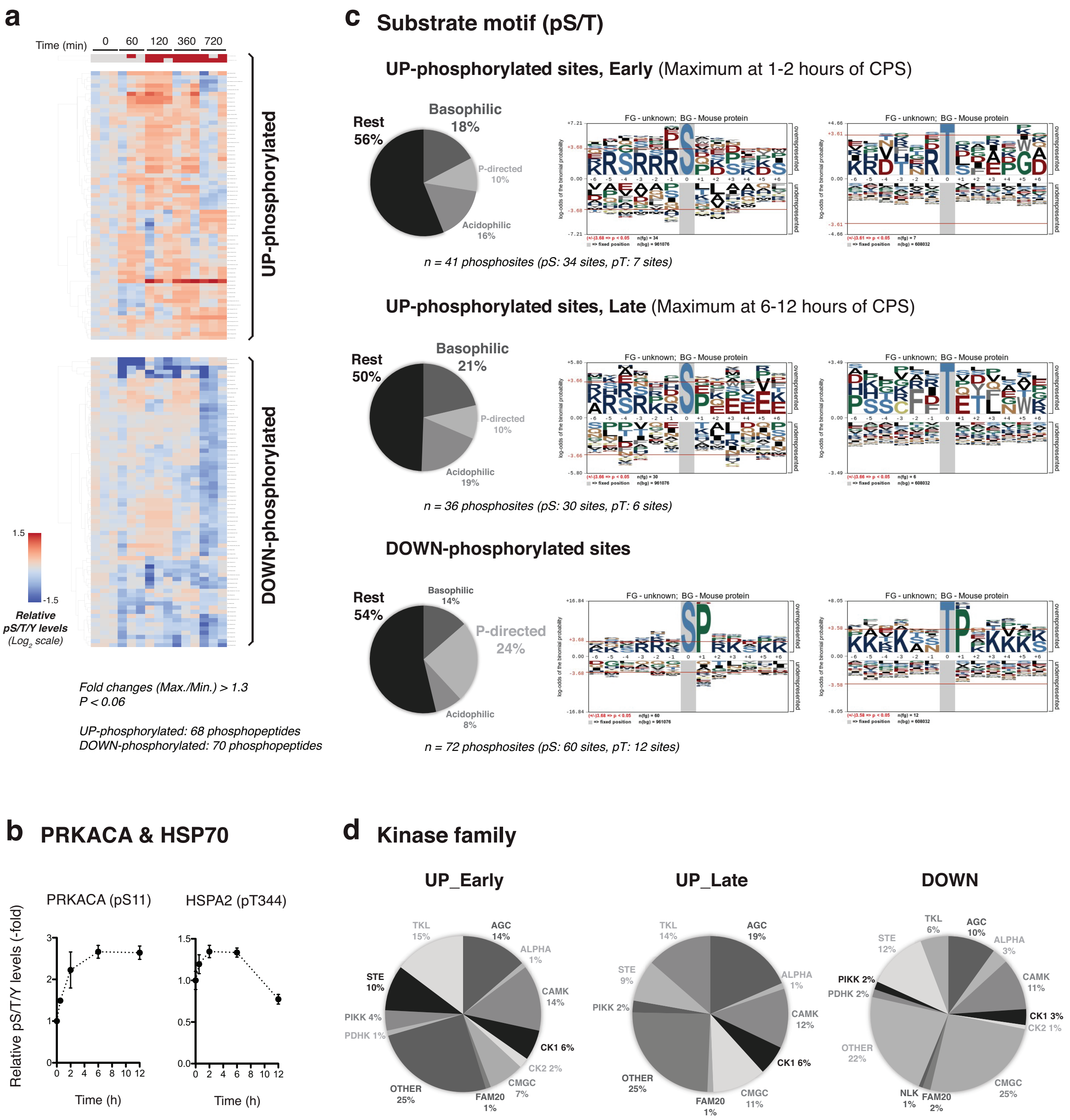


**Supplementary Fig. 4 | Dynamic protein phosphorylation in AML12 cells following DNAJB1/PRKACA generation by CPS. a,** Cluster map of the proteomics dataset representing the time-resolved phosphorylation combined from three biological replicates. Data processing with the stringency filters described in Supplementary Fig. 2b results in 138 representative dynamics over the 12-hour time period. 'UP-phosphorylated' proteins corresponded to at least a 30% increase compared to the average phosphorylation levels at time zero (Max./0 hr value > 1.3-fold, n = 68). 'DOWN-phosphorylated' proteins corresponded to at least a 50% decrease from maximum to minimum and a 20% or more decrease from time zero to minimum average phosphorylation levels within the 12-hour timeslot (0 hr/Min. value > 1.2-fold & Max./Min. value > 1.5-fold, n = 70). **b,** Fold-change in phosphorylation of PRKACA and HSP70 at indicative times. Phosphorylation levels at each time point are normalized to the values at 0 hours (*n*=3. Mean with s.e.m). **c,** Pie charts (left) summarizing clustered phosphorylation-site motifs (predicted by The Kinase Library[43]) and enriched sequence motifs (right) derived from serine/threonine phosphorylation sites sorted by their dynamics. 'UP_Early' protein modification sites showed maximum phosphorylation at 0.5-2 hours of CPS, and 'UP_Late' member sites had maximum phosphorylation at 6-12 hours of CPS. 'DOWN' indicates the modification sites of the 'Down-phosphorylated' proteins in (**a**). **d,** Pie charts summarizing analysis of the three phosphorylation groups (UP_Early, UP_Late, DOWN) against the clustered human serine/threonine kinome (predicted by The Kinase Library[43]).

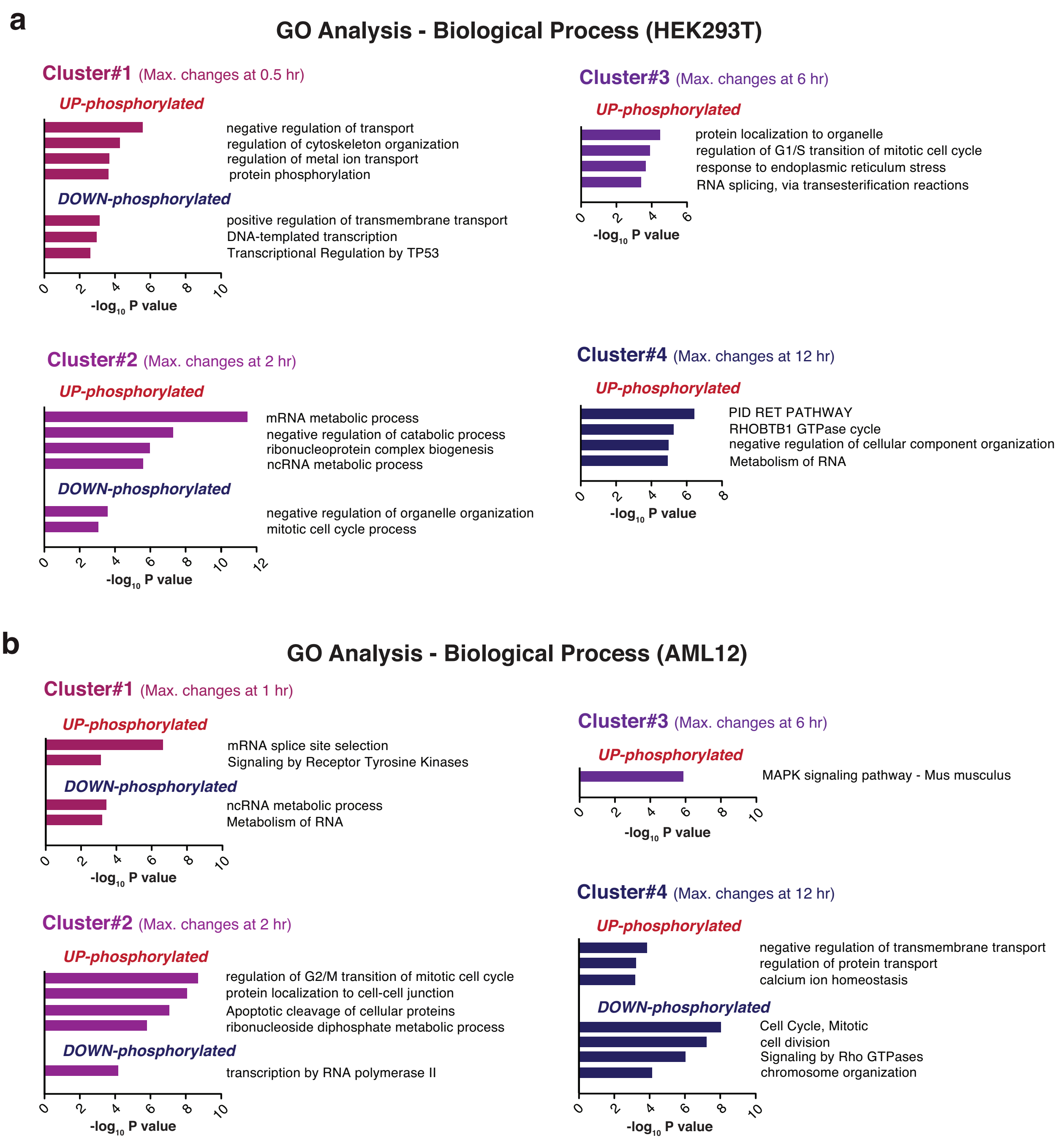


**Supplementary Fig. 5 | Gene ontology analysis. a, b,** GO analysis of each cluster in HEK293T (**a**) and AML12 (**b**) cells.

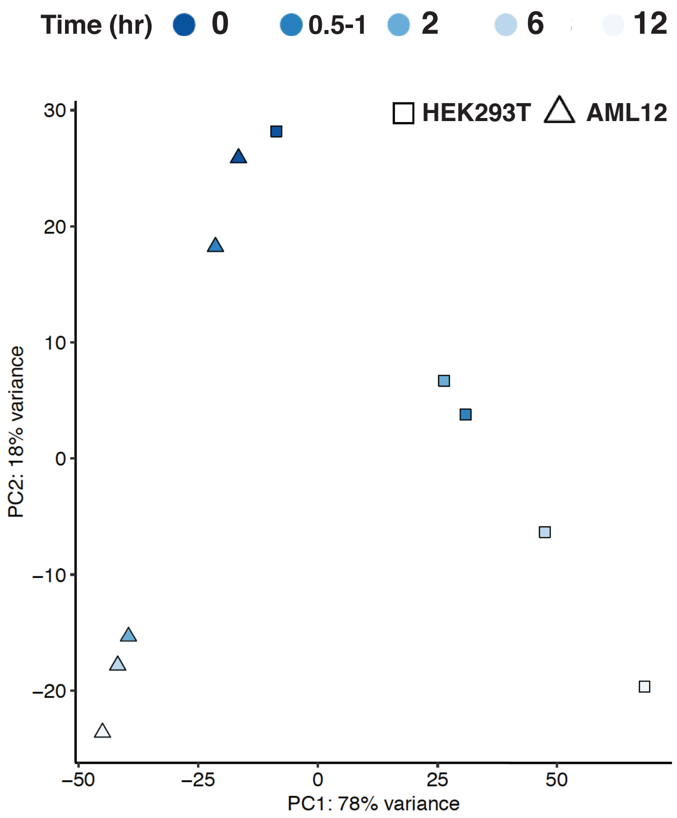


**Supplementary Fig. 6 | Post-translational generation of DNAJB1/PRKACA leads to cell-specific changes in protein phosphorylation levels.** Principal component analysis (PCA) of the dynamic phosphorylation changes in HEK293T (n = 801, satisfying $P < 0.06$ in Supplementary Fig. 2b) and AML12 (n = 602, satisfying $P < 0.06$ in Supplementary Fig. 2b) cells following generation of DNAJB1/PRKACA by CPS. To ensure a fair comparison of phosphorylation dynamics between two datasets, without any bias toward fold-changes, we specifically chose the group [b] from Supplementary Fig. 2b, c as the input.

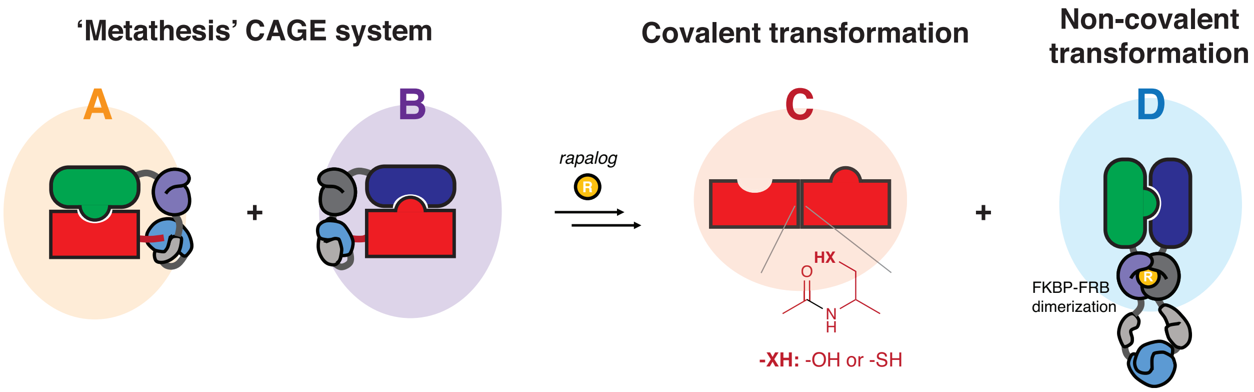


**Supplementary Fig. 7 | Protein 'metathesis' using the proximity induced CPS.** Protein trans-splicing results in the simultaneous exchange of pairs of function molecules attached to the CAGE modules via covalent and non-covalent interactions. This 'A + B → C + D' transformation can be thought of as a type of protein 'metathesis'.
